# *Ta*ExpA6, *VRT-A2* and *Ta*GW2 genes differentially affect grain weight, grain number and yield of wheat through their physiological determinants

**DOI:** 10.64898/2025.12.04.692342

**Authors:** Lucas Vicentin, Daniel F. Calderini

**Author notes:** Author for correspondence: (D.F. Calderini).

## Abstract

Physiological basis of the trade-off between grain number (GN) and thousand grain weight (TGW) is key to improve wheat yield. To that end, three wheat line groups were assessed at conventional (300-350 pl m^-2^) and low (44 pl m^-2^) plant rates in the field: *Ta*ExpA6 (ectopic expression), *Ta*GW2 (*TaGW2* triple mutant), and *Ta*P1xGW2A (*VRT-A2* ectopic expression and *TaGW2-A* knock-out), together with their wild-types (WT). Except *Ta*P1xGW2A, the manipulated lines increased TGW over their WTs. However, only the transgenic *TaExpA6* line reached higher grain yield (GY) than its WT, whereas increased TGW recorded by the triple knockout of *TaGW2* was fully compensated by reduced spike number (SpN) and grain number per spike (GNS). Contrasting effects of *Ta*ExpA6 and *Ta*GW2 lines were found on the ovary weight and floret dynamics, associated with the trade-off between GW and GN. Interestingly, the overexpression of *TaExpA6* gene acts in grain tissues at post-anthesis avoiding the overlap with GN determination, while *TaGW2* disruption constrains tillering, likely as a pleiotropic effect, and shifts intra-spike resource allocation promoting early ovary growth at the expense of distal floret development and GNS. Low plant rate increased resource allocation to the spike, mitigating the GW–GNS trade-off and increasing both traits simultaneously.

## 1. Introduction

Wheat (*T. aestivum*) is a key crop to ensure global food security as it provides ∼20% of calories and protein to the human diet (Braun *et al*., 2010). Current average rate of wheat yield increase of ∼0.9% p.a. is below the minimum required to keep pace with the estimated 50% increase in wheat production needed by 2050 to deal with a projected human population of 9.7 billion people (Ray *et al*., 2013; Fischer *et al*., 2014; FAO, 2017; Vollset *et al*., 2020; van Dijk *et al*., 2021), imposing a daunting challenge for agriculture.

Historically, grain yield of wheat has been improved by increasing grain number per unit area (GN) through plant breeding and crop management (Calderini *et al*., 1995; Reynolds *et al*., 2009; Slafer *et al*., 2021). Over the last years, several studies, most of them from a molecular perspective, have focused on grain weight (GW) as a way to improve wheat yield (Kumar *et al*., 2016; Simmonds *et al*., 2016; Brinton *et al*., 2017; Wang *et al*., 2019; Calderini *et al*., 2021; Guo *et al*., 2022). A recent meta-analysis proposed that selecting for both increased GN and GW is the most promising strategy to further improve harvest index and grain yield of wheat (Xi *et al*., 2024). However, the trade-off between average grain weight (TGW) and GN is a bottleneck for improving GY as demonstrated across a wide range of wheat genotypes and environments (Bustos *et al*., 2013; Slafer *et al*., 2014; Quintero *et al*., 2018; Molero *et al*., 2019; Rivera-Amado *et al*., 2019).

The trade-off between TGW and GN became evident in attempts to increase wheat yield by improving GW (Slafer *et al*., 2021), and this negative compensation was manifested when GW was either increased by conventional breeding or targeted genetic modifications using molecular tools (Wiersma *et al*., 2001; Okamoto and Takumi, 2013; Brinton *et al*., 2017; Wang *et al*., 2018; Zhai *et al*., 2018; Adamski *et al*., 2021; Jablonski *et al*., 2021; Milner *et al*., 2021; Mora-Ramirez *et al*., 2021; Brunner *et al*., 2024). Different genes have been successful to increase TGW. Among them, the long-glume P1 locus from *T. polonicum*, codified by the *VRT-A2* gene, has been studied due to its positive effect on GW through the modulation of grain length (Okamoto and Takumi, 2013; Adamski *et al*., 2021; Liu *et al*., 2021). Wheat NILs ectopically expressing the *VRT-A2* gene increased TGW by 5.5% relative to its WT but did not impact on GY due to a concomitant reduction of GN (Adamski *et al*., 2021). In a similar manner, the most successful enhancement of GW was achieved by the knockout of the gene *TaGW2*, switching off its negative control of GW and allowing a very high increase of TGW over the WT, i.e. ∼20% (Wang *et al*., 2018). Other genetic studies have additionally pointed out the potential of switching off the *TaGW2* gene to increase TGW of wheat (Su *et al*., 2011; Yang *et al*., 2012; Qin *et al*., 2014; Sestili *et al*., 2019), however, these attempts have failed to increase GY of wheat up to date due to the negative impact of *TaGW2* knockout on grain number (GN) (Zhai *et al*., 2018; Kis *et al*., 2024; Vicentin *et al*., 2024).

Regardless of its relevance for yield improvement, the underlying genetic and physiological processes involved in the trade-off between TGW and GN remain mostly unknown. Therefore, the actual impact of *VRT-A2* and *GW2* genes on GN and the dissection of their effect on GN subcomponents is a prerequisite to the use of these genes in plant breeding. On the other hand, the proof of concept that the trade-off between GW and GN could be broken has been provided by Calderini *et al*. (2021) and Vicentin *et al*. (2024) by transgenic wheat lines overexpressing the *TaExpA6* gene in grains from 10 days after anthesis (DAA) on. The higher TGW conferred by the

*TaExpA6* gene resulted in into improved GY (+11%) over the segregant WT in field experiments, without a trade-off affecting GN or any other agronomic trait (Calderini *et al*., 2021). Regarding that it is widely accepted that wheat is not limited by the source of assimilates during the effective grain filling period (Slafer and Savin, 1994; Richards, 1996; Dreccer *et al*., 1997; Borrás *et al*., 2004; Calderini *et al*., 2006; Serrago *et al*., 2013; Borrill *et al*., 2015; Lichthardt *et al*., 2020; Slafer *et al*., 2021; Slafer *et al*., 2023), the GN-GW trade-off must be attributed to other non-competitive causes (Acreche and Slafer, 2006).

In addition to specific genes, simple crop management decisions such as plant rate also modify TGW in different extent, offering a complementary opportunity to assess the mechanisms involved in grain weight determination. However, contradictory responses have been found for GN and TGW when contrasting plant rates were evaluated. For instance, TGW enhancements between 6 and 17% have been reported in response to plant rate reductions ranging from 40 to 85% (Ellen, 1990; Li *et al*., 2016; Hasan *et al*., 2022). On the contrary, in the experiments carried out by Lloveras *et al*. (2004) and Bustos *et al*. (2013), plant rate reductions within the same range affected GN (from -18 to 13%) but with almost negligible effect on TGW, which differs from the aforementioned increases reported for this key trait. Therefore, the underlying causes of plant rate impact on either GN or GW, but especially on GW, are still poorly understood. Speculatively, it has been proposed that the response of GW to plant rate reductions would be associated with changes in biomass partitioning and light quality conditions (Red:Far red ratio) within the canopy (Ugarte *et al*., 2010; Dreccer *et al*., 2022; Hasan *et al*., 2022).

Given the significance of the trade-off between GN and GW and the lack of knowledge about its causes in wheat, the objectives of the present study were (i) to evaluate the response of grain yield, GN, GW and their interaction in unique genotypes showing contrasting trade-off between GW and GN, (ii) to quantify the effect of low plant rate on grain yield components and the trade-off between GW and GN, and (iii) to integrate tiller, floret dynamics and GW assessment to uncover physiological mechanisms accounting for the negative interaction between both major yield components of wheat.

## 2. Materials and methods

### 2.1 Field conditions and experimental setup

Three field experiments were performed in a Typic Hapludand soil at the Experimental Station of Universidad Austral de Chile (EEAA) in Valdivia (39°47’S, 73°14’W) during three growing seasons: 2021-2022 (Exp. 1), 2022-2023 (Exp. 2) and 2023-2024 (Exp. 3). In the experiments three pairs of wheat genotypes were assessed: (i) line *Ta*ExpA6, a transgenic line with ectopic expression of the α-expansin gene *TaExpA6* in grains, as described by Calderini *et al*. (2021) and its segregant wild type cv. Fielder; (ii) line *Ta*P1xGW2A, a wheat NIL of cv. Paragon with the combined effect of *T. polonicum VRT-A2* gene and *TaGW2-A* loss-of-function mutation, and its segregant wild type cv.

Paragon; and (iii) line *Ta*GW2, a triple mutant of the gene *TaGW2* and its segregant WT. Each pair of genotypes is referred to as the ExpA6, P1xGW2A and GW2 groups, respectively, and were chosen due to their improved TGW and contrasting effect on GN when they were assessed under field conditions. Lines *Ta*P1xGW2A and *Ta*GW2 evidenced a trade-off between GW and GN in former studies (Okamoto and Takumi, 2013; Wang *et al*., 2018; Zhai *et al*., 2018; Adamski *et al*., 2021), whereas the line *Ta*ExpA6 enhances GW without a negative impact on GN (Calderini *et al*., 2021). The *Ta*ExpA6 line carries a construct of *TaExpA6* gene (REFSEQ v.1.1: *TraesCS4A02G034200*), controlled by the wheat *puroindoline-b* (*PinB*) gene promoter (REFSEQ v.1.1: *TraesCS7B02G431200*), which allows its expression in the endosperm, aleurone, and pericarp tissues of developing grains (Gautier *et al*., 1994; Digeon *et al*., 1999). The line *Ta*P1xGW2A carries both the ectopic expression of *VRT-A2* gene (long-glume phenotype of *T. polonicum*, Adamski *et al*., 2021) and a mutation in the A-genome homeologue of *TaGW2*, detailed in Simmonds *et al*. (2016). Unlike the line *Ta*P1xGW2A, line *Ta*GW2 carries mutations in all three homeologous copies of the *TaGW2* gene, resulting in a non-functional protein. A detailed characterization of the mutated alleles was previously documented by Simmonds *et al*. (2016) and Wang *et al*. (2018).

All experiments were set in a randomized complete block design with four replications. In experiment 1 (Exp. 1), lines of the ExpA6 and P1xGW2A groups were sown on September 21^st^, 2021 at two plant rates: conventional plant rate of 300 plants m^−2^ (CPR300) and low plant rate of 44 plants m^−2^ (LPR), the last under a squared arrangement, i.e. 0.15 × 0.15 m. In experiment 2 (Exp. 2), the lines *Ta*P1xGW2A and its WT were left aside, since no difference in TGW was observed between them in Exp. 1. These lines were replaced by the GW2 lines. Hence, four lines were sown in Exp. 2: (i) line *Ta*ExpA6, (ii) its segregant WT Fielder, (iii) line *Ta*GW2 and (iv) its segregant WT. In Exp. 2, GW2 and ExpA6 groups were sown at both CPR300 (300 plants m^−2^) and LPR (44 plants m^−2^). In order to match the flowering dates of both groups of genotypes, GW2 lines with longer-crop cycle were sown earlier on August 20^th^, 2022, while the ExpA6 lines were sown later, on September 2^nd^ 2022. In experiment 3 (Exp. 3), as the different sowing date did not have effect, both the ExpA6 and GW2 lines were sown on September 4^th^, 2023, but under a single plant density of 350 plants m^−2^ (CPR350).

Across experiments, each plot was 2 m long and 1.2 m wide, consisting of 9 rows spaced 0.15 m apart. Optimal management practices were carried out to all plots to keep the crop free of any biotic and abiotic constraints. Supplementary drip irrigation was applied in all experiments, complementing rainfall, to prevent water shortage until physiological maturity. The experimental sites were treated with 4 t ha⁻¹ of CaCO₃ one month before sowing to prevent potential aluminium toxicity from low soil pH. Fertilizers were applied at sowing to avoid nutritional shortage with rates of 150 kg N ha⁻¹, 150 kg P₂O₅ ha⁻¹, and 100 kg K₂O ha⁻¹, and an additional 150 kg N ha⁻¹ was applied at mid-tillering. All pest and diseases were chemically prevented or controlled following the manufacturer treatment recommendations. Weather conditions (i.e. air temperature and incident solar radiation) were recorded hourly throughout the crop cycle at the EEAA’s meteorological station (http://agromet.inia.cl/) located by 150 m from the experimental plots across experiments.

### 2.2 Crop and floret development and carpel growth dynamics

Crop phenology was recorded according to the decimal code scale (Zadoks *et al*., 1974) twice a week along the crop cycle, registering each stage when 50% of the main shoots reached each developmental stage. Thermal time was calculated as the summation of daily average temperature [(T_max_+T_min_)/2] and a base temperature of 0°C was used.

From one week before booting to 7 DAA, four spikes of similar size and development were selected and harvested from the main stratus (i.e. main shoots plus primary tillers with similar size and development) of each experimental unit twice a week to register the developmental stage of each floret (floral score) from the central spikelet.

For this assessment, the scale proposed by Waddington *et al*. (1983) was used, as illustrated in Ferrante *et al*. (2013). Florets were numbered following the same procedure described by González *et al*. (2003), namely from F1 to Fn regarding their position from the spike rachis, being F1 the floret primordium closest to the rachis, and Fn the most distal primordium in each spikelet. At pollination (i.e., Waddington stage 10; W10), the weight of ovaries from florets at positions F1 to F5 (if present) was measured by sampling 20 ovaries of each floret position as previously (Vicentin *et al*., 2024). To test the association between ovary weight at pollination and final grain weight, 10 spikes of similar size and development from the main stratus were sampled from each plot at harvest to quantify grain weight at each grain position from central spikelets of the spike (spike mapping), following the same procedure as Vicentin *et al*. (2024).

In experiments 2 and 3, the ovary growth dynamics was monitored during the overlapping window between GN and GW determination, i.e. between booting and a week after anthesis (Calderini *et al*., 2021). To this end, 12 florets of the F2 position from the 4 central spikelets of three spikes were additionally sampled following the same procedure to remove the ovary at booting (Z45) and heading (Z55). Ovaries were dried in an oven at 65°C for 48 h to record the ovary dry weight using an electronic balance (Mettler Toledo, XP205DR, Greifensee, Switzerland) as previously (Hasan *et al*., 2011). Ovary growth rate from booting to W10 was estimated as the difference between ovary dry weight at W10 and booting, divided by the thermal time elapsed between these samplings. In Exp. 3, 10 spikes per plot were harvested at Z65 to determine the number of fertile florets at anthesis. To that end, all florets from half the spikelets (i.e. all the spikelets along one longitudinal side of the spike), including the apical spikelet, were considered. A floret was recorded as fertile when it reached stage 10, according to the scale defined by Waddington *et al*. (1983). All counted florets, except those from the apical spikelet, were multiplied by two to estimate the total number of fertile florets per spike.

### 2.3 Crop sampling and measurements

In Exps. 2 and 3, the number of tillers was counted once a week in four labelled plants per plot from Z21 up to one week after anthesis (Z65) as in Evers *et al*. (2006). The duration of tiller appearance (Dur_Ap_), duration of tiller senescence (Dur_Sen_), tillering rate (T_rate_) and tiller mortality (T_mort_) were calculated for each plot as:

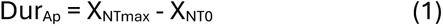

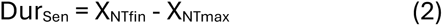

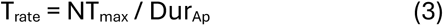

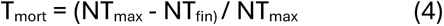

where: X_NT0_, X_NTmax_ and X_NTfin_ represent the timing (in days after emergence) of Z21, maximum tiller number and final tiller number, respectively; NT_max_ and NT_fin_ represent the maximum and final number of tillers per plant recorded at each plot.

Biomass samples were harvested from each plot at both anthesis (Z65) and physiological maturity, i.e. when grain dry matter levelled off as in Hasan *et al*. (2011). The sampled plants were harvested from central rows, leaving rows acting as border to protect the sampling area. In CPD plots, biomass samples were 0.5 m long at anthesis and 1.0 m at maturity. In LPD plots, samples of 0.9 m long (six plants) were taken at each harvest time as in Hasan *et al*. (2022).

Biomass samples were divided into main shoots and tillers (except in the anthesis samples of Exp.1), and then separated into blade leaves, stems plus sheath leaves and spikes. At maturity, plant height was measured from the ground to the base of the spike in main stems of 10 and 6 plants per plot of CPR and LPR treatments, respectively. After drying for 48 h at 60°C, biomass samples were weighed using an electronic balance (WTC2000, Radwag, Radom, Poland) to determine total above-ground biomass (AGB). To assess potential variations in stem water-soluble carbohydrates (WSC) accumulation at anthesis, a representative sample was taken from main stems at this phenological stage in CPR plots of Exps. 2 and 3 for analysis of WSC content. WSC were determined by the anthrone method, following the protocol proposed by McDonald and Henderson (1964). At maturity, grains were threshed and separated from other spike structures (chaff) using a laboratory thresher (LD 180, Wintersteiger, Austria). After that, grain samples were oven-dried at 60°C for 48 h and weighed to record grain yield, using the same electronic balance than for biomass samples. Harvest index (HI) was calculated as the ratio between grain yield and above-ground biomass. GN was obtained by counting all the grains from each sample with a seed counter (Contador 2, Pfeuffer, Kitzingen, Germany), and this trait was expressed as number per square metre (GN) by dividing the sampled number by the surface area. Average GW was calculated by dividing total grain biomass by grain number and expressed as thousand grain weight (TGW) in grams.

### 2.5 Statistical analysis

The effects of genotype, plant rate and their interaction on yield and associated traits, ovary weight, floret development and tillering parameters were assessed by analysis of variance (ANOVA), with genotype and plant rate as fixed factors. Data from each experiment was analysed separately. Differences across the recorded variable data were considered statistically significant at a probability level of 5% using Fisheŕs least squares mean differences (LSD) test. ANOVA was performed using InfoStat software, version 2020 (Di Rienzo *et al*., 2020). Linear regression analyses were performed to test the fit of data, slopes and associations between variables using the least sum-of-squares fitting method in GraphPad Prism 8 software. Extra sum of squares F test was used to identify significant differences in the slopes and origins of the regressions between genotype groups and plant rate treatments.

## 3. Results

### 3.1 Crop phenology and climate conditions

Across all three experiments, the *TaExpA6* overexpressing line and its WT counterpart (Fielder) exhibited a shorter crop cycle than the P1xGW2A and GW2 groups, as plotted in Fig. 1. In Exp. 1, the whole crop cycle of ExpA6 and P1xGW2A groups in the CPR treatment lasted 116 and 129 days, respectively (200°Cd difference). At the same plant rate, and across Exps. 2 and 3, the ExpA6 and GW2 groups reached physiological maturity in 125 and 141 days, respectively, showing a similar difference (i.e. 222°Cd) as observed for the P1xGW2A group in Exp. 1. As both last groups were developed from the cultivar Paragon, their longer crop cycle, especially the emergence-booting phase (Fig. 1), can be ascribed to the Paragon genetic background. Likely, lines developed from Paragon have higher photoperiod sensitivity than Fielder, the genetic background of the overexpressed ExpA6 line (see photoperiod in Table 1). On the other hand, as expected, no phenological difference was found between the modified lines and their respective WTs of each group, Regarding plant rate, a significant (p<0.05), but narrow effect, was found for this treatment on crop phenology (Table S1), as the Bo-An period was 2 days longer in the LPR than under the CPR treatment in P1xGW2A lines, and 1 d longer in ExpA6 and GW2 lines. Additionally, the grain filling period was 3.5 days longer in LPR treatments, averaged across the three groups of genotypes in Exps. 1 and 2.

**Figure 1.**
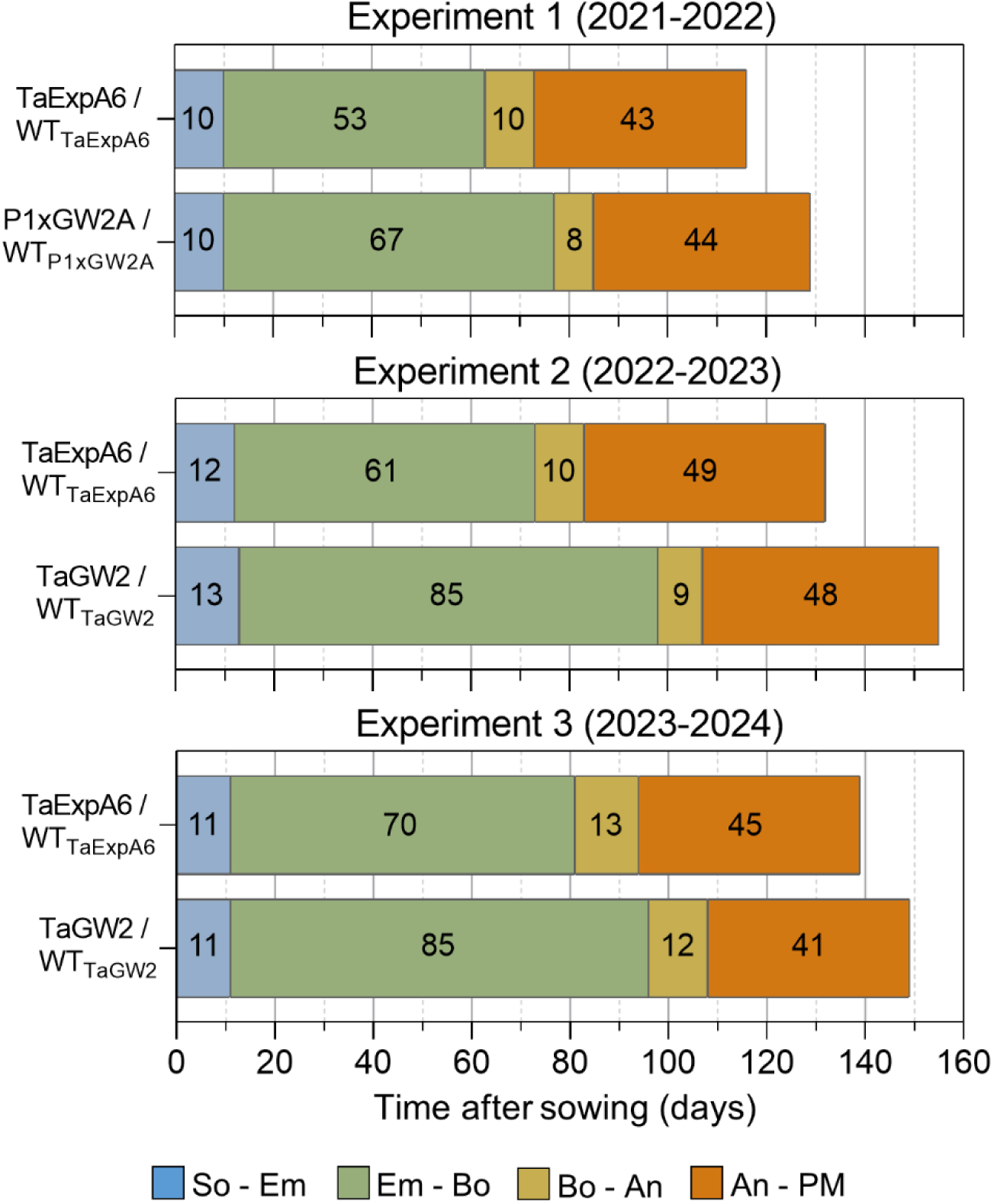
Crop phenology of ExpA6, P1xGW2A and GW2 groups recorded at CPR plots in experiments 1, 2 and 3. Given the absence of phenological difference, each modified line and its respective WT within each group are depicted in the same bar. Numbers denote the mean duration of each phenological phase.

**Table 1.**
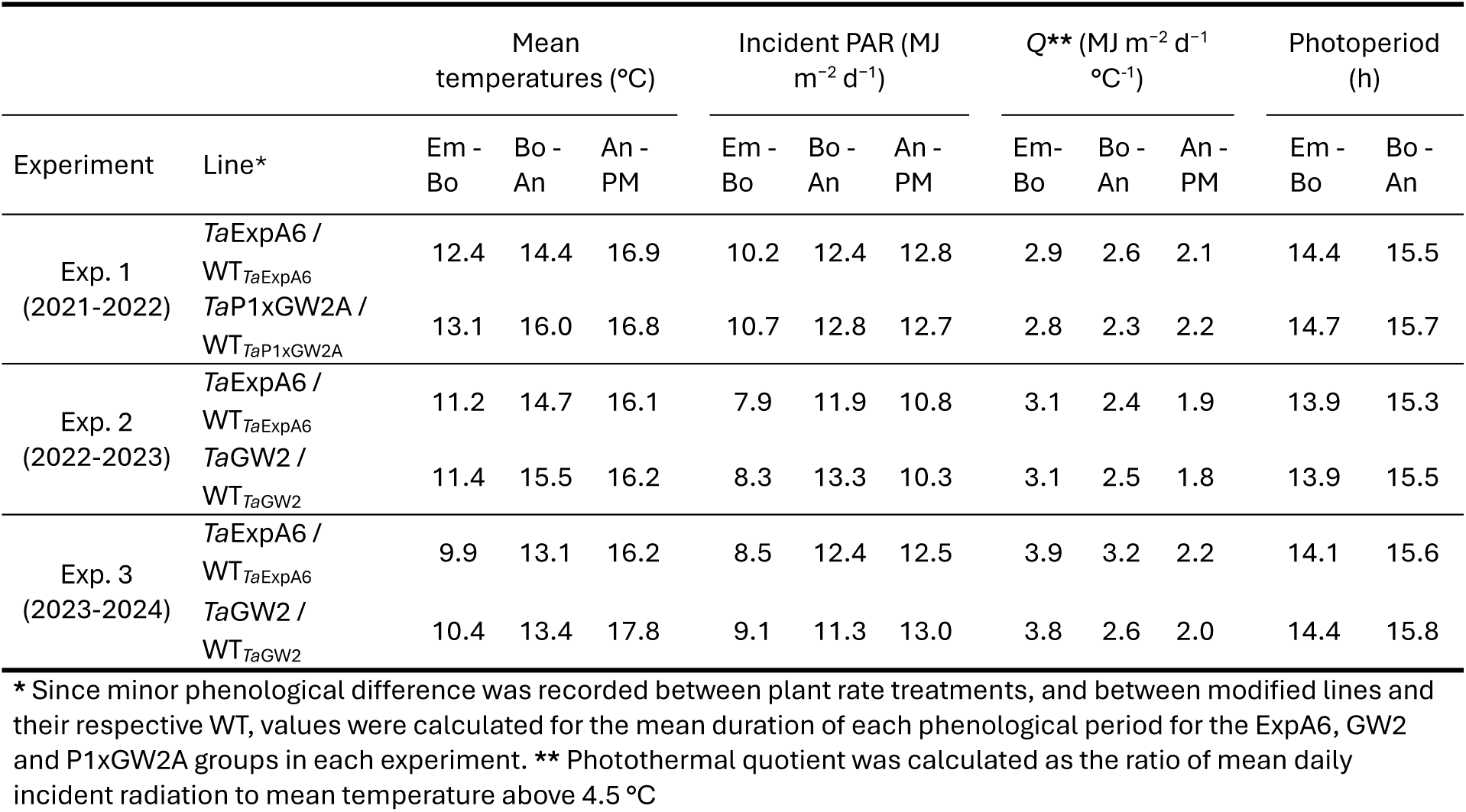
Average mean temperature, Incident photosynthetically active radiation (PAR), Photothermal quotient (*Q*) and Photoperiod during the Emergence-Booting (Em-Bo), Booting-Anthesis (Bo-An) and Anthesis-Physiological Maturity (An-PM) periods for each group of lines during Experiments 1, 2 and 3.

Climate conditions during the experiments were in accordance with the southern Chile environment, showing favourable temperature and high photothermal quotient for temperate crops. Average temperatures across genotypes between emergence and anthesis (Em-An) were 13.9°C (Exp. 1), 13.2°C (Exp. 2) and 11.7°C (Exp. 3), whereas mean temperatures during grain filling were 16.9°C, 16.2°C and 17°C in Exps. 1, 2 and 3, respectively (Table 1). In line with the lower temperature recorded up to anthesis in Exp. 3, the photothermal quotient (*Q*) in this experiment was 24% higher during Em-An than in the previous experiments (i.e. 3.4 *vs.* 2.7 MJ m^-2^ day^-1^ °C^-1^, the last averaged across Exps. 1 and 2). During grain filling, *Q* ranged from 1.9 MJ m^-2^ day^-1^ °C^-1^ (Exp. 2) to 2.2 MJ m^-2^ day^-1^ °C^-1^ (Exp. 1). Despite the observed phenological difference between groups, genotypes within each experiment were exposed to similar weather conditions (Table 1). Across Exps. 1 and 2, differences in weather exposure between plant rate treatments were negligible (Table S2), in agreement with the minimal phenological differences observed in response to plant rate reduction.

### 3.2 Grain yield across experiments and treatments

A broad range of grain yield (GY) was recorded across the experiments, spanning from 833 to 1443 g m^-2^ (Table 2), with the highest grain yield reached in Exp. 3, in agreement with the utmost *Q* value during the critical period for grain number determination (Table 1). Despite this variation, similar GY was achieved among the genotype groups through the experiments. For instance, average grain yield of each group at CPR300 across experiments were 1050 g m^-2^ in ExpA6, 1099 g m^-2^ in P1xGW2A and 1048 g m^-2^ in GW2 group (Table 2). On the other hand, the contribution of main stems (MS) and tillers to total GY differed among groups at CPR. While similar contribution between both stem categories was recorded in the ExpA6 group, GY of MS was proportionally higher than tillers in P1xGW2A and GW2 groups, i.e. 60 and 75% of total GY, respectively (Table S3), which is likely due to genetic background differences between the parental cultivars, i.e. Fielder and Paragon.

**Table 2.**
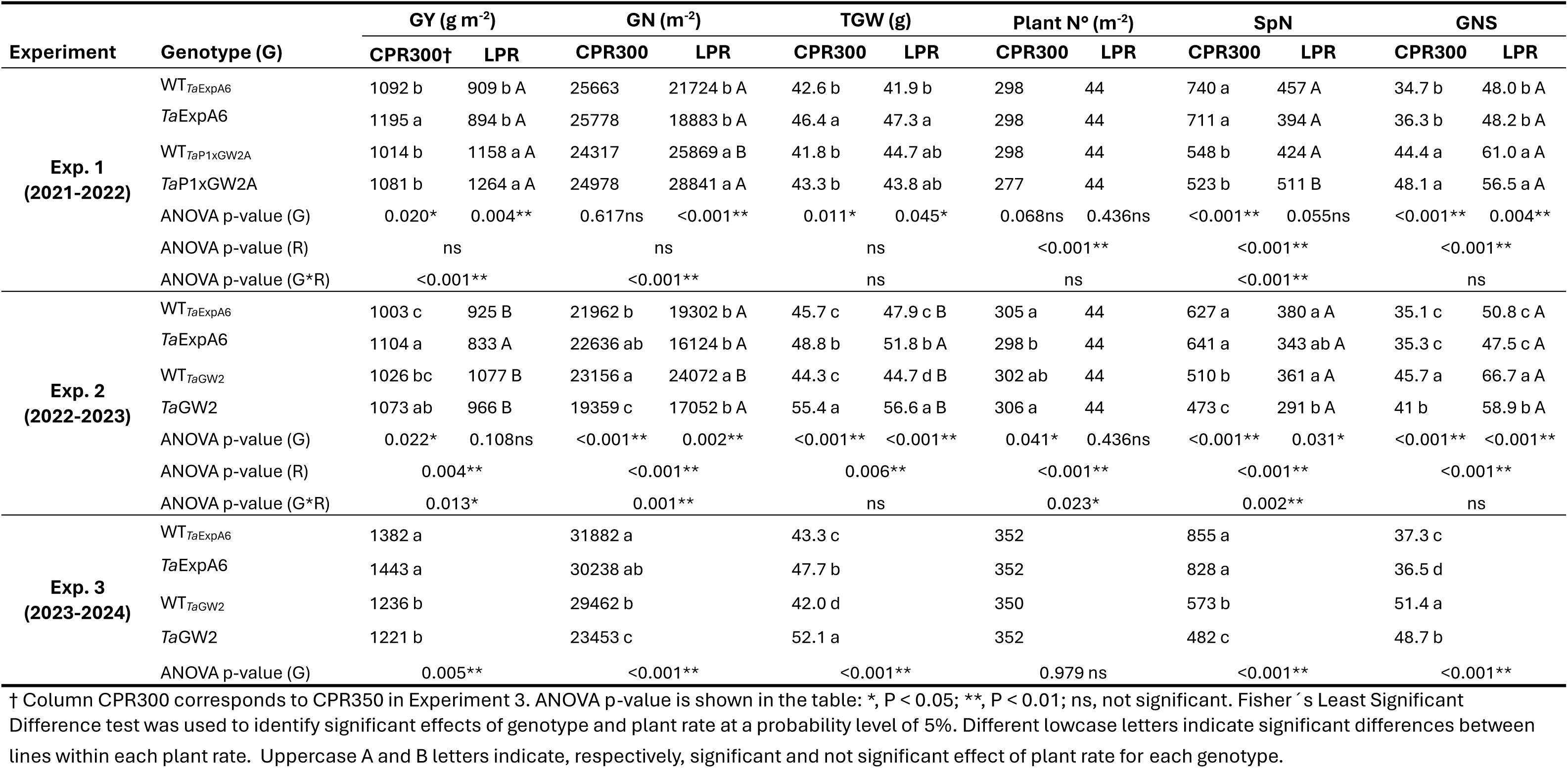
Grain yield (GY), grain number per square metre (GN), thousand grain weight (TGW), plant number per square metre, spike number per square metre (SpN) , and grain number per spike (GNS) recorded in ExpA6, P1xGW2A and GW2 groups in Experiments 1, 2 and 3

Within each group, the *Ta*ExpA6 line outyielded (p<0.05) its WT under farmers plant rate (CPR300) by 9.4 and 10.1% in Exps. 1 and 2, respectively (Table 2). In Exp. 3, GY of line TaExpA6 was only 4.4% higher than its WT, which was not statistically different (p>0.05; Table 2). In the P1xGW2A group, solely sown in Exp. 1, there was not GY differences (p> 0.05) between the *Ta*P1xGW2A line and its WT (Table 2). Likewise, similar GY (p>0.05) were achieved by the triple mutant line *Ta*GW2 and its WT at CPR treatment in Exps. 2 and 3, i.e. +4.6% and -1.2%, respectively. In the LPR treatment, similar (p>0.05) GY was attained by each modified line and its respective WT in the three genotype groups (Table 2).

Regarding the effect of plant rate, no differences in GY were observed when the contrasting plant rates were averaged across genotypes in Exps. 1 and 2 (i.e. 1074 and 1003 g m^-2^ at CPR300 and LPR, respectively). However, a significant interaction between genotype and plant rate was found for GY in these experiments (Table 2).

When the modified lines and their WTs were averaged within each genotype group, GY was negatively (p<0.05) affected by the plant rate reduction in the ExpA6 lines (-21%), no effect (p>0.05) of LPR was found in the GW2 lines, and GY was increased (p<0.05) under LPR in P1xGW2A lines (+15.6%) (Table 2).

Across genotypes, plant rate treatments and experiments, GY was positively and closely associated with AGB (Fig. 2A). However, no association was found between GY and HI (Fig. 2B). Accordingly, the latter trait was highly conservative between the modified lines and their WTs (Table 3). Under LPR condition, a subtle increase (p<0.05) of HI was found, i.e. an average increase of 3-4 percentage points over CPR across genotypes and experiments.

**Figure 2.**
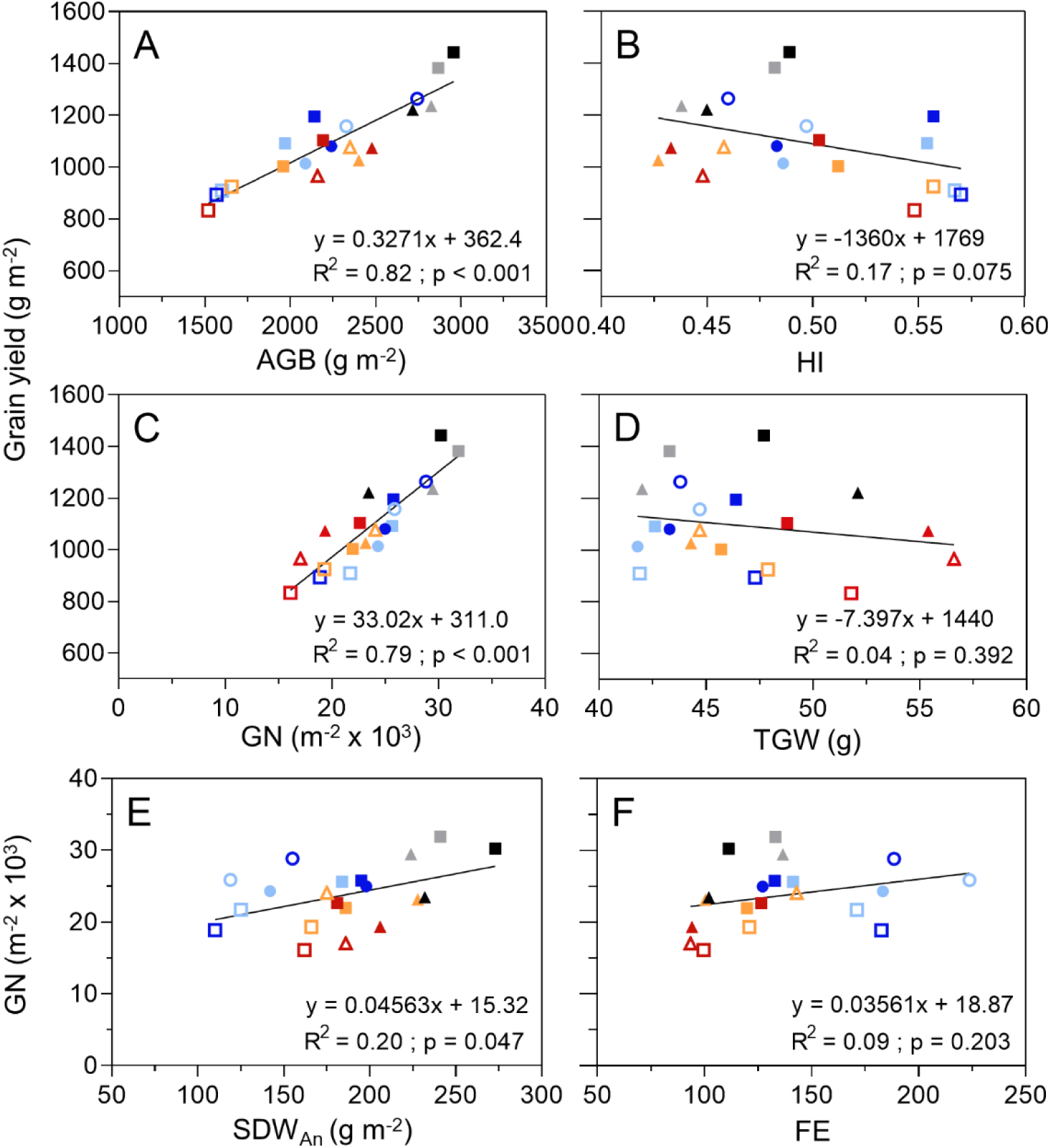
Linear relationship between grain yield and (A) total aboveground biomass, (B) harvest index, (C) grain number per square metre and (D) average grain weight. Linear relationship between grain number per square metre and (E) spike dry weight at anthesis and (F) fruiting efficiency. Lines ExpA6 (squares), P1xGW2A (circles) and GW2 (triangles) sown at CPR (closed symbols) and LPR (open symbols). Dark and lighter coloured symbols represent modified (Mod) and WT lines within each group, respectively. Data from experiments 1, 2 and 3 are represented, respectively, in blue (Mod)/light blue (WT), red (Mod)/orange (WT), and black (Mod)/grey (WT).

**Table 3.** Total aboveground biomass (AGB), harvest index (HI), plant height, spike dry weight at anthesis (SDW_An_), fruiting efficiency (FE) and stem water-soluble carbohydrates content at anthesis (WSC_An_) recorded in ExpA6, P1xGW2A and GW2 groups in Experiments 1, 2 and 3.

| Experiment | Genotype (G) | AGB (g m<sup>-2</sup>) |  | HI |  | Height (cm) |  | SDW<sub>An</sub> (g m<sup>-2</sup>) |  | FE (GN SpikeDM<sub>Ant</sub><sup>-1</sup>) |  | Stem WSC<sub>An</sub> |  |
| --- | --- | --- | --- | --- | --- | --- | --- | --- | --- | --- | --- | --- | --- |
|  |  | CPR300† | LPR | CPR300 | LPR | CPR300 | LPR | CPR300 | LPR | CPR300 | LPR | CPR300 | LPR |
| Exp. 1<br>(2021-2022) | WT<sub>TaExpA6</sub> | 1970 | 1601 c A | 0.55 a | 0.57 a A | 79.9 c | 63.6 d A | 184 | 125 A | 141 | 171 A |  |  |
|  | TaExpA6 | 2143 | 1569 c A | 0.56 a | 0.57 a A | 82.1 c | 69.9 c A | 195 | 110 A | 133 | 183 A |  |  |
|  | WT<sub>TaP1xGW2A</sub> | 2090 | 2328 b A | 0.49 b | 0.50 b A | 88.1 b | 89.6 b B | 142 | 119 A | 183 | 224 A |  |  |
|  | TaP1xGW2A | 2241 | 2744 a A | 0.48 b | 0.46 c B | 94.1 a | 95.6 a B | 198 | 155 A | 127 | 188 A |  |  |
|  | ANOVA p-value (G) | 0.135ns | <0.001** | <0.001** | <0.001** | <0.001** | <0.001** | 0.146ns | 0.156ns | 0.185ns | 0.527ns |  |  |
|  | ANOVA p-value (R) | ns |  | ns |  | <0.001** |  | <0.001** |  | 0.011* |  |  |  |
|  | ANOVA p-value (G*R) | <0.001** |  | 0.046* |  | <0.001** |  | ns |  | ns |  |  |  |
| Exp. 2<br>(2022-2023) | WT<sub>TaExpA6</sub> | 1959 a | 1658 b A | 0.51 a | 0.56 a A | 87.9 c | 80.4 c A | 186 b | 166 A | 120 ab | 121 ab B | 191,9 a |  |
|  | TaExpA6 | 2193 b | 1520 b A | 0.50 a | 0.55 a A | 93.2 b | 85.7 b A | 181 b | 162 A | 127 a | 100 b A | 195,9 a |  |
|  | WT<sub>TaGW2</sub> | 2402 c | 2351 a B | 0.43 b | 0.46 b A | 100.7 a | 96.8 a A | 228 a | 175 A | 100 bc | 143 a A | 73,2 b |  |
|  | TaGW2 | 2478 c | 2160 a A | 0.43 b | 0.45 b A | 100.1 a | 95.7 a A | 206 ab | 186 A | 94 c | 94 b B | 84,3 b |  |
|  | ANOVA p-value (G) | <0.001** | 0.003** | <0.001** | <0.001** | <0.001** | <0.001** | 0.043* | 0.711ns | 0.017* | 0.036* | <0.001** |  |
|  | ANOVA p-value (R) | <0.001** |  | <0.001** |  | <0.001** |  | 0.008** |  | ns |  |  |  |
|  | ANOVA p-value (G*R) | 0.017* |  | ns |  | ns |  | ns |  | 0.006** |  |  |  |
| Exp. 3<br>(2023-2024) | WT<sub>TaExpA6</sub> | 2868 |  | 0.48 a |  | 100.4 b |  | 241.0 |  | 133.0 |  | 177,5 a |  |
|  | TaExpA6 | 2958 |  | 0.49 a |  | 108.4 a |  | 273.0 |  | 111.0 |  | 159,0 a |  |
|  | WT<sub>TaGW2</sub> | 2826 |  | 0.44 b |  | 110.9 a |  | 224.0 |  | 137.0 |  | 76,8 b |  |
|  | TaGW2 | 2717 |  | 0.45 b |  | 110.1 a |  | 232.0 |  | 102.0 |  | 79,6 b |  |
|  | ANOVA p-value (G) | 0.384 ns |  | <0.001** |  | <0.001** |  | 0.056 ns |  | 0.145 ns |  | 0.013* |  |

† Column CPR300 corresponds to CPR350 in Experiment 3. ANOVA p-value is shown in the table: \*, P < 0.05; \*\*, P < 0.01; ns, not significant. Fisher's Least Significant Difference test was used to identify significant effects of genotype and plant rate at a probability level of 5%. Different low case letters indicate significant differences between lines within each plant rate. Uppercase A and B letters indicate, respectively, significant and not significant effect of plant rate for each genotype.

### 3.3 Genotype and plant rate effect on average grain weight

TGW was distinctively affected by the genotypes. Contrary to the expected, the *Ta*P1xGW2A line reached similar (p>0.05) TGW than its WT in Exp. 1 (Table 2). By contrast, both the transgenic *Ta*ExpA6 and the *Ta*GW2 triple mutant lines increased (p<0.05) TGW over their WTs, though at different magnitudes. The overexpression of the *Ta*ExpA6 gene improved TGW above its WT by 9, 6.8 and 10.2% at CPR in Exps. 1, 2 and 3, respectively (Table 2), while the triple mutant line reached a 25 and 24% higher TGW than its WT in Exps. 2 and 3, respectively, under CPR (Table 2). Under the LPR treatment, line *Ta*ExpA6 also improved TGW over its WT, but as expected, this increase was higher in relative terms than in the CPR treatment (i.e. +13% and +8.1% in Exp. 1 and 2, respectively; Table 2). On the other hand, line *Ta*GW2 improved (p<0.05) TGW over its WT by 26.7% when it was sown at LPR, which was similar to the increase recorded in the CPR treatment. The positive effect of the *TaExpA6* construct and *TaGW2* mutation on TGW was observed in both stems categories, i.e. MS and tillers across experiments and plant rate treatments (Table S3). The positive impact of the LPR treatment on TGW was found when MS and tillers were analysed separately, whereas a negligible effect was found for overall TGW (Table 2). Thus, across all lines, TGW at LPR was 8.8% and 6.4% higher in MS (p<0.05), and 5.5% and 7.5% higher in tillers (p<0.05) than that of CPR in Exps. 1 and 2, respectively (Table S3).

### 3.4 Individual GW was associated to ovary growth up to anthesis

When final grain weight of grain positions G1 to G5 from the central spikelets of spikes were plotted against their ovary weight at pollination (W10 stage, Waddington *et al*. scale), a positive linear association was found between both traits across lines, grain/floret positions and experiments (R^2^>0.79; p<0.001; Fig. 3A-D). Across all spikelets and grain positions, individual GW increased by a mean of 12.3% and 20.1% in the *Ta*ExpA6 and *Ta*GW2 lines, respectively, relative to their corresponding WTs, when averaged across experiments. Likewise, individual GW across all positions was, on average, 19.9% and 18.9% higher under the LPR treatment than under the CPR treatment in Experiments 1 and 2, respectively.

**Figure 3.**
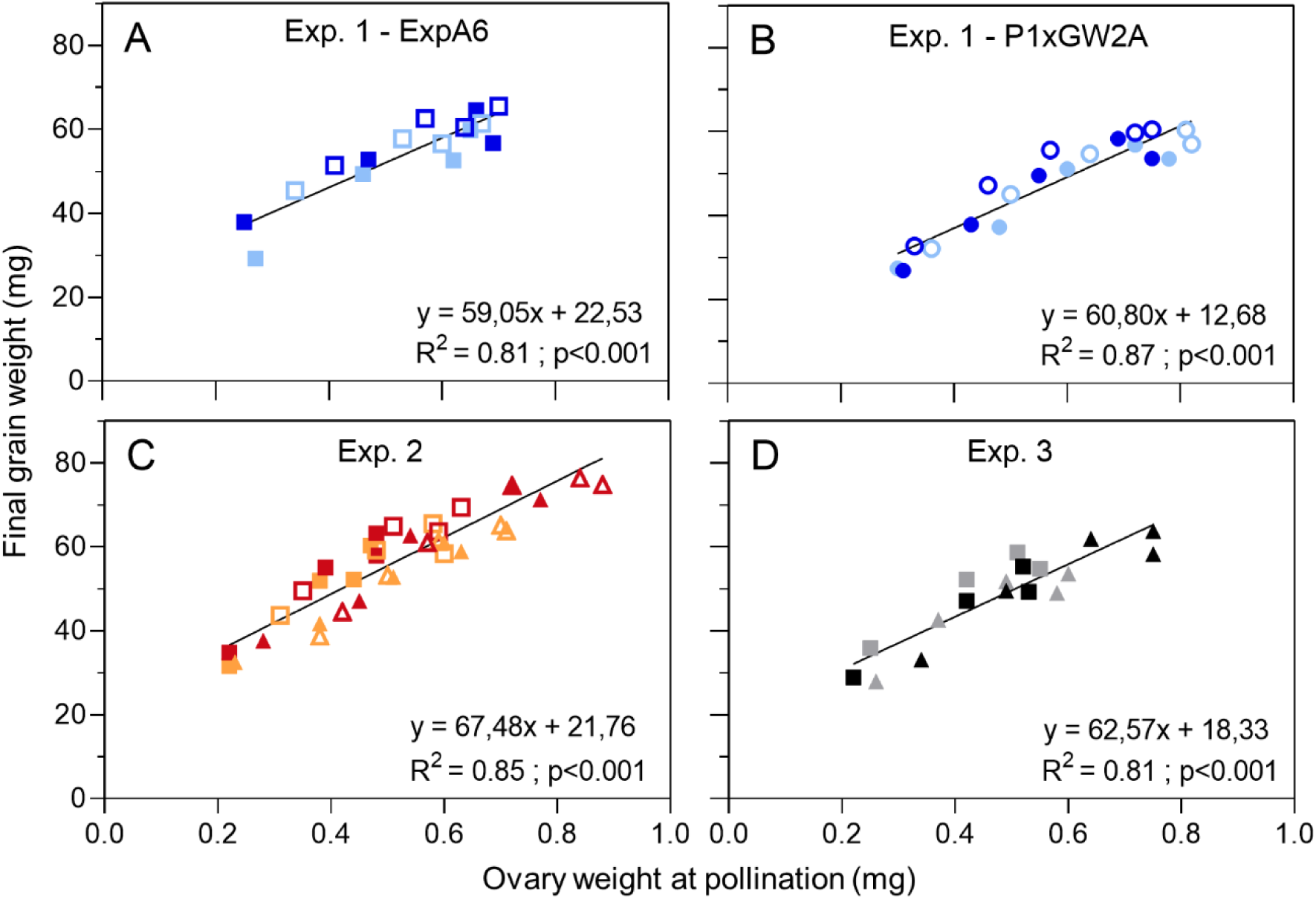
Relationship between final grain weight and ovary weight at pollination (W10, Waddington et al., 1983) of grain positions G1 to G5 (if present) from the central spikelets of the spike of lines ExpA6 (squares), P1xGW2A (circles) and GW2 (triangles) sown at CPR (closed symbols) and LPR (open symbols). Dark and lighter coloured symbols represent modified (Mod) and WT lines within each group, respectively. Data from experiments 1, 2 and 3 are represented, respectively, in blue (Mod)/light blue (WT), red (Mod)/orange (WT), and black (Mod)/grey (WT). Data of lines from (A) ExpA6 and (B) P1xGW2A groups in Exp. 1, and (C) all lines in Exp. 2 and (D) Exp. 3 were analysed separately due to significant differences in the slopes and origin of the equations, according to extra sum of squares F test (Table S5).

Additionally, the dynamics of ovary weight of floret F2 was recorded to have a deeper insight on the time-course of the ovary growth. To this aim, the ovary weight from central spikelets were measured at booting (Bo), heading (Hd) and pollination (W10) in the ExpA6 and GW2 groups under both CPR and LPR treatments in Exps. 2 and 3.

Interestingly, the ovary weight of line *Ta*ExpA6 and its WT did not differ (p>0.05) at any of the three sampling times under both CPR and LPR treatments in Exps. 2 and 3 (Fig. 4A, C). On the other hand, the F2 ovary of line *Ta*GW2 was heavier (p<0.05) than that of its WT as early as Bo across experiments and plant rates. This difference, ranging between 20 and 31% (p<0.05), was maintained up to W10 (Fig. 4B, D). Remarkably, the LPR treatment increased (p<0.05) the ovary weight over the CPR300 treatments from Bo onwards in ExpA6 lines, and from Hd onwards in GW2 lines (Fig. 4A, B), resulting in 24 and 17% heavier ovaries at pollination, respectively.

**Figure 4.**
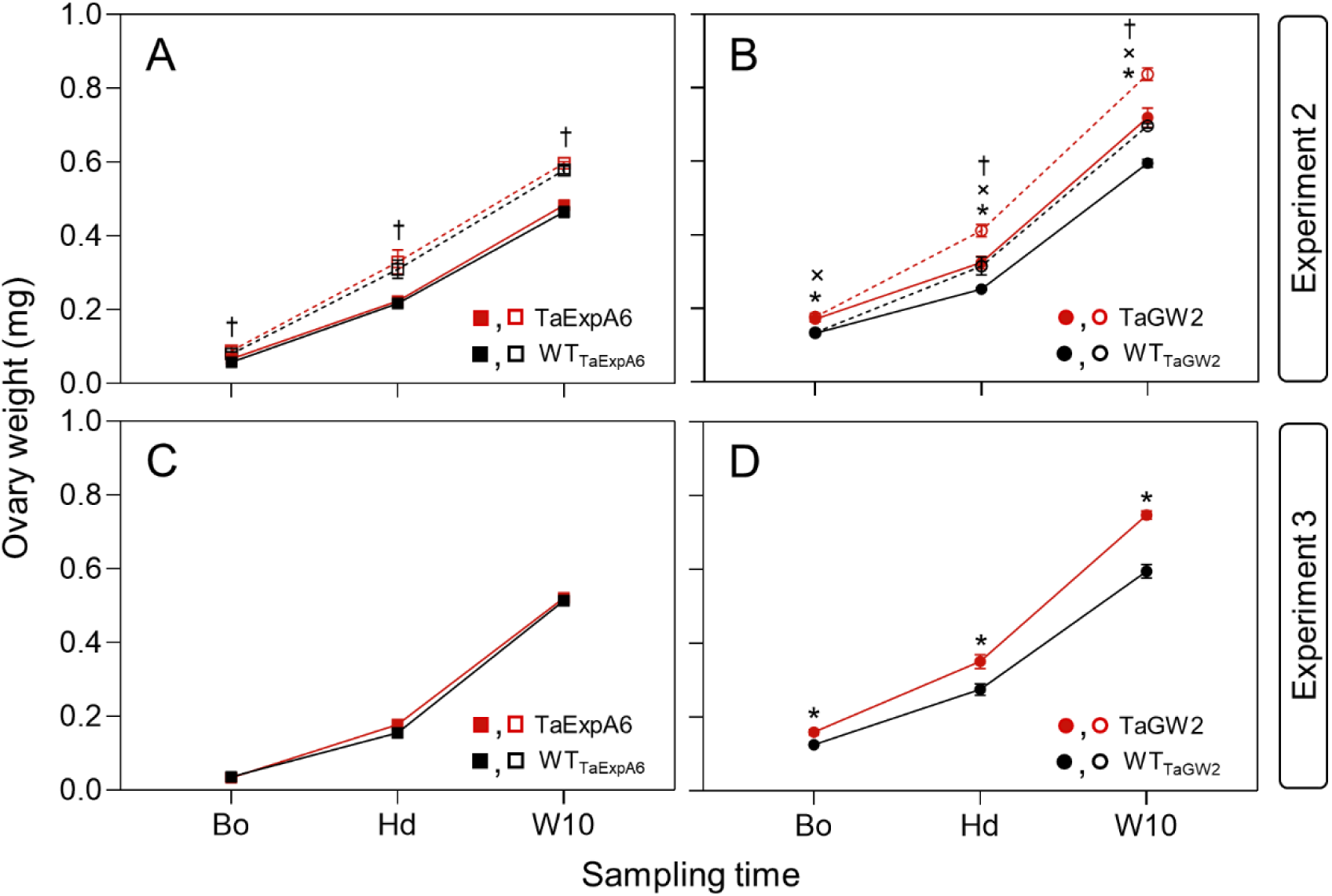
Weight of ovaries from floret F2 of central spikelets of main stem spikes at booting (Bo), heading (Hd) and pollination (W10) in experiments 2 and 3, corresponding to the ExpA6 lines (A, C) and GW2 lines (B, D). Closed symbols and solid lines represent CPR treatments in experiments 2 and 3. Open symbols and dashed lines represent LPR treatments in experiment 2. A factorial two-way ANOVA was performed separately for each group of lines, with genotype and plant rate as fixed effects. No significant interaction was found between factors. Significant effect of genotype on ovary weight at CPR and LPR treatments is denoted by **\*** and **^x^** symbols, respectively. Significant effect of plant rate is denoted by † symbol.

Differences in ovary weight between line *Ta*GW2 and its WT were due to the ovary growth rate during the Bo-W10 period, which was 14.7% higher (p<0.05) in the triple mutant line averaged across experiments and plant rates (Table 4; Fig. S1A).

**Table 4.** Individual spike dry weight at anthesis (SDW_An_) and fruiting efficiency (FE_ind_) in Experiments 1, 2 and 3. Duration of the booting-pollination (Bo-W10) period and growth rate of F2 ovaries from central spikelets of main stems of ExpA6 and GW2 groups in Experiments 2 and 3.

| Experiment | Genotype / Plant rate | Individual $SDW_{An}$ | | $FE_{ind}$ | | Duration of Bo-W10 period ($^{\circ}Cd$) | | Ovary growth rate ($mg\ ^{\circ}Cd \times 10^{-2}$) | |
| --- | --- | --- | --- | --- | --- | --- | --- | --- | --- |
|  |  | CPR300† | LPR | CPR300 | LPR | CPR300 | LPR | CPR300 | LPR |
| Exp. 1<br>(2021-2022) | WT<sub>TaExpA6</sub> | 0.31 | 0.52 | 113.5 | 98.1 |  |  |  |  |
|  | TaExpA6 | 0.33 | 0.45 | 108.1 | 108.5 |  |  |  |  |
|  | WT<sub>TaP1xGW2A</sub> | 0.26 | 0.36 | 185.6 | 176 |  |  |  |  |
|  | TaP1xGW2A | 0.35 | 0.37 | 139.2 | 153.1 |  |  |  |  |
|  | WT<sub>TaExpA6</sub> | 0.41 |  | 105.8 b |  |  |  |  |  |
|  | TaExpA6 | 0.39 |  | 108.3 b |  |  |  |  |  |
|  | WT<sub>TaP1xGW2A</sub> | 0.31 |  | 180.8 a |  |  |  |  |  |
|  | TaP1xGW2A | 0.36 |  | 146.2 ab |  |  |  |  |  |
|  | **Dif. (ExpA6)** | **-4.9%** |  | **2.4%** |  |  |  |  |  |
|  | **Dif. (P1xGW2A)** | **16.1%** |  | **-19.1%** |  |  |  |  |  |
|  | CPR300 | 0.31 b |  | 136.6 |  |  |  |  |  |
|  | LPR | 0.43 a |  | 133.9 |  |  |  |  |  |
|  | **Dif. (Plant rate)** | **38.7%** |  | **-2.0%** |  |  |  |  |  |
|  | ANOVA p-value (G) | 0.074 |  | 0.008 |  |  |  |  |  |
|  | ANOVA p-value (R) | <0.001 |  | 0.857 |  |  |  |  |  |
|  | ANOVA p-value (G*R) | 0.128 |  | 0.897 |  |  |  |  |  |
| Exp. 2<br>(2022-2023) | WT<sub>TaExpA6</sub> | 0.27 e | 0.42 c | 128.3 ab | 120.4 b | 144.0 | 166.8 | 0.28 | 0.30 |
|  | TaExpA6 | 0.28 e | 0.50 b | 125.0 b | 94.6 c | 140.2 | 168.3 | 0.30 | 0.30 |
|  | WT<sub>TaGW2</sub> | 0.35 d | 0.47 bc | 132.6 ab | 144.9 a | 131.4 | 145.2 | 0.35 | 0.39 |
|  | TaGW2 | 0.44 bc | 0.67 a | 92.8 c | 88.3 c | 137.3 | 153.3 | 0.40 | 0.43 |
|  | WT<sub>TaExpA6</sub> | 0.35 c |  | 124.4 b |  | 155.4 a |  | 0.29 c |  |
|  | TaExpA6 | 0.39 b |  | 109.9 c |  | 154.2 a |  | 0.30 c |  |
|  | WT<sub>TaGW2</sub> | 0.41 b |  | 138.7 a |  | 138.3 c |  | 0.37 b |  |
|  | TaGW2 | 0.56 a |  | 90.6 d |  | 145.3 b |  | 0.41 a |  |
|  | **Dif. (ExpA6)** | **11.4%** |  | **-11.7%** |  | **-0.8%** |  | **3.4%** |  |
|  | **Dif. (GW2)** | **36.6%** |  | **-34.7%** |  | **5.1%** |  | **10.8%** |  |
|  | CPD300 | 0.34 b |  | 119.7 |  | 138.2 b |  | 0.33 b |  |
|  | LPD | 0.51 a |  | 112.1 |  | 158.4 a |  | 0.36 a |  |
|  | **Dif. (Plant rate)** | **50.0%** |  | **-6.3%** |  | **14.6%** |  | **9.1%** |  |
|  | ANOVA p-value (G) | <0.001 |  | <0.001 |  | <0.001 ** |  | <0.001 ** |  |
|  | ANOVA p-value (R) | <0.001 |  | 0.081 |  | <0.001 ** |  | 0.046 * |  |
|  | ANOVA p-value (G*R) | 0.038 |  | 0.014 |  | 0.062 ns |  | 0.597 ns |  |

**Table 4.** Continued
| Experiment | Genotype / Plant rate | Individual SDW<sub>An</sub> |  | FE<sub>ind</sub> |  | Duration of Bo-W10 period (°Cd) |  | Ovary growth rate (mg °Cd x 10<sup>-2</sup>) |  |
| --- | --- | --- | --- | --- | --- | --- | --- | --- | --- |
|  |  | CPR300† | LPR | CPR300 | LPR | CPR300 | LPR | CPR300 | LPR |
| Exp. 3<br>(2023-2024) | WT<sub>TaExpA6</sub> | 0.31 b |  | 121.1 b |  | 166 |  | 0.29 b |  |
|  | TaExpA6 | 0.33 b |  | 112.2 b |  | 160.6 |  | 0.30 b |  |
|  | WT<sub>TaGW2</sub> | 0.36 b |  | 142.9 a |  | 154.5 |  | 0.31 b |  |
|  | TaGW2 | 0.43 a |  | 119.2 b |  | 160 |  | 0.37 a |  |
|  | **Dif. (ExpA6)** | **6.5%** |  | **-7.3%** |  | **-3.3%** |  | **3.4%** |  |
|  | **Dif. (GW2)** | **19.4%** |  | **-16.6%** |  | **3.6%** |  | **19.4%** |  |
|  | ANOVA p-value (G) | 0.007 |  | 0.019 |  | 0.151 ns |  | 0.002 ** |  |

† Column CPR300 corresponds to CPR350 in Experiment 3. The percentage difference between each modified line and its respective WT, and between plant rates is written in bold. ANOVA p-value is shown in the table: \*, P < 0.05; \*\*, P < 0.01; ns, not significant. Fisher's Least Significant Difference test was used to identify significant effects of genotype and plant rate at a probability level of 5%. Different lowercase letters indicate significant differences (p<0.05).

Alternatively, the positive effect of low plant rate on ovary weight can be attributed to both the duration of the Bo-W10 period, and the higher rate of ovary growth between these stages relative to the CPR300 treatment, i.e. a 15 and 9% increase under LPR, respectively (Table 4; Fig. S1C, D).

In addition to ovary weight, the content of water-soluble carbohydrates in stems at anthesis (WSC_An_) was also recorded in CPR plots of Exps. 2 and 3, given that they represent a key source of assimilates for grain filling. However, no association (R^2^ = 0.03, p> 0.05) was found between TGW and WSC_An_ (Fig. S2).

### 3.5 Genotype and plant rate effect on grain number and its numerical and physiological determinants

Despite the successful increase of TGW accomplished by both the transgenic and triple mutant lines over their WTs, the regression analysis showed that differences in GY were mainly explained by grain number (GN) and not by TGW across experiments (Fig. 2C and D). The P1xGW2A lines reached similar (p>0.05) GN at conventional and low plant rates in Exp. 1, in addition to the lack of effect of the *Ta*P1xGW2A line on TGW (Table 2). Likewise, the *Ta*ExpA6 construct had no effect (p>0.05) on GN at the CPR treatment across experiments (Table 2), explaining the improvement of GY achieved by the ectopic expression of the *Ta*ExpA6 gene in growing grains. In Exp. 3, although GN did not show differences (p>0.05) between the transgenic *Ta*ExpA6 line and its WT at CPR, this trait was 5.2% lower in the modified line than the WT, apparently due to low minimum temperatures (i.e. from 0.6 °C to 1.3°C recorded at meteorological station, 1.65 m above the ground) between booting and anthesis in the overexpressing line (Fig. S3), affecting the expected superior GY of this line as in Exps. 1 and 2. No statistical differences (p> 0.05) in GN were found between the ExpA6 lines under the LPR treatment, but line *Ta*ExpA6 also showed a lower value of GN relative to the WT (-14.7% when averaging Exps. 1 and 2; Table 2). In contrast, TGW increase recorded in line *Ta*GW2 was always offset by the reduction (p<0.05) of GN relative to its WT at the CPR treatment in Exps. 2 and 3 (-16.4 and -20.4%, respectively) and also under LPR (-29%) in Exp. 2 (Table 2).

The effect of treatments on GN was analysed by its numerical subcomponents, i.e. the spike number per square meter (SpN) and grain number per spike (GNS). GN sub-components were not affected by either the *Ta*P1xGW2A or the *Ta*ExpA6 line compared with their respective WTs (Table 2; Table S3). On the other side, the *TaGW2* triple mutant line showed lower GN than its WT across the experiments due to a clear decrease (p<0.05) of both SpN and GNS relative to its WT (Table 2; Table S3).

Specifically, SpN in the *Ta*GW2 line was decreased by 7.4% (CPR; Exp. 2), 16% (CPR; Exp. 3), and 19.5% (LPR; Exp. 2) relative to its WT, while GNS was decreased by 10%, 5.3%, and 11.7%, respectively. Regarding the effect of LPR, a genotype and plant rate interaction (p<0.05) was found for GN, mainly due to a differential response of SpN under LPR treatment (Table 2), whereas GNS was substantially improved (p<0.05) by plant rate reduction across all lines, i.e. +36.5% (Table 2).

Surprisingly, no associations (p>0.05) between GN and either SDW_An_ or FE (Fig. 2E, F) were found in this study. Moreover, neither SDW_An_ nor FE showed differences between the modified lines and their respective WTs (Table 3). SDW_An_ and FE were further analysed in MS at individual spike level (i.e. SDW_An_ and FE per spike - FE_Ind_ -). Across groups, the only difference (p<0.05) was found in line *Ta*GW2, which increased individual SDW_An_ by 30.5% relative to its WT, but on the contrary FE_Ind_ decreased by 28.6% across Exps. 2 and 3 (Table 4). LPR treatment also increased (p<0.05) individual SDW_An_ in Exps. 1 and 2, but in a higher magnitude (i.e. +44.6%), and without any change (p>0.05) in FE_Ind_ across lines and experiments (Table 4). Accordingly, the increase of individual SDW_An_ by genotypic effect was fully compensated by reductions in FE_Ind_ (Fig. S4A), whereas the negative response showed by FE_Ind_ was proportionally lower than the individual SDW_An_ increase under LPR (Fig. S4B).

### 3.6 Tillering capacity of genotypes explained the response of SpN in the assessed lines

The time-course of tillering was measured in Exps. 2 and 3, given its importance to explain SpN, and in turn, GN. Very contrasting tillering behaviours were found between the manipulated lines and their WTs, as well as between plant rate treatments. In the ExpA6 group, closely similar (p>0.05) tillering dynamics were observed between both lines when sown at CPR in Exps. 2 and 3 (Fig. 5A, E; Table 5). On the contrary, the triple mutant line *Ta*GW2 developed a lower (p<0.05) number of tillers and SpN than its WT across plant rate treatments and experiments (Fig 5B, D, F; Table 5). This was evidenced by NT_max_ and NT_fin,_ which dropped by 31 and 23%, respectively, relative to its WT (Fig 5B, D, F; Table 5). At LPR, line *Ta*ExpA6 developed a lower number of maximum (NT_max_) and final (NT_fin_) tillers per plant than its WT, though this decrease was not statistically (p> 0.05) significant (Fig 5C; Table 5).

**Figure 5.**
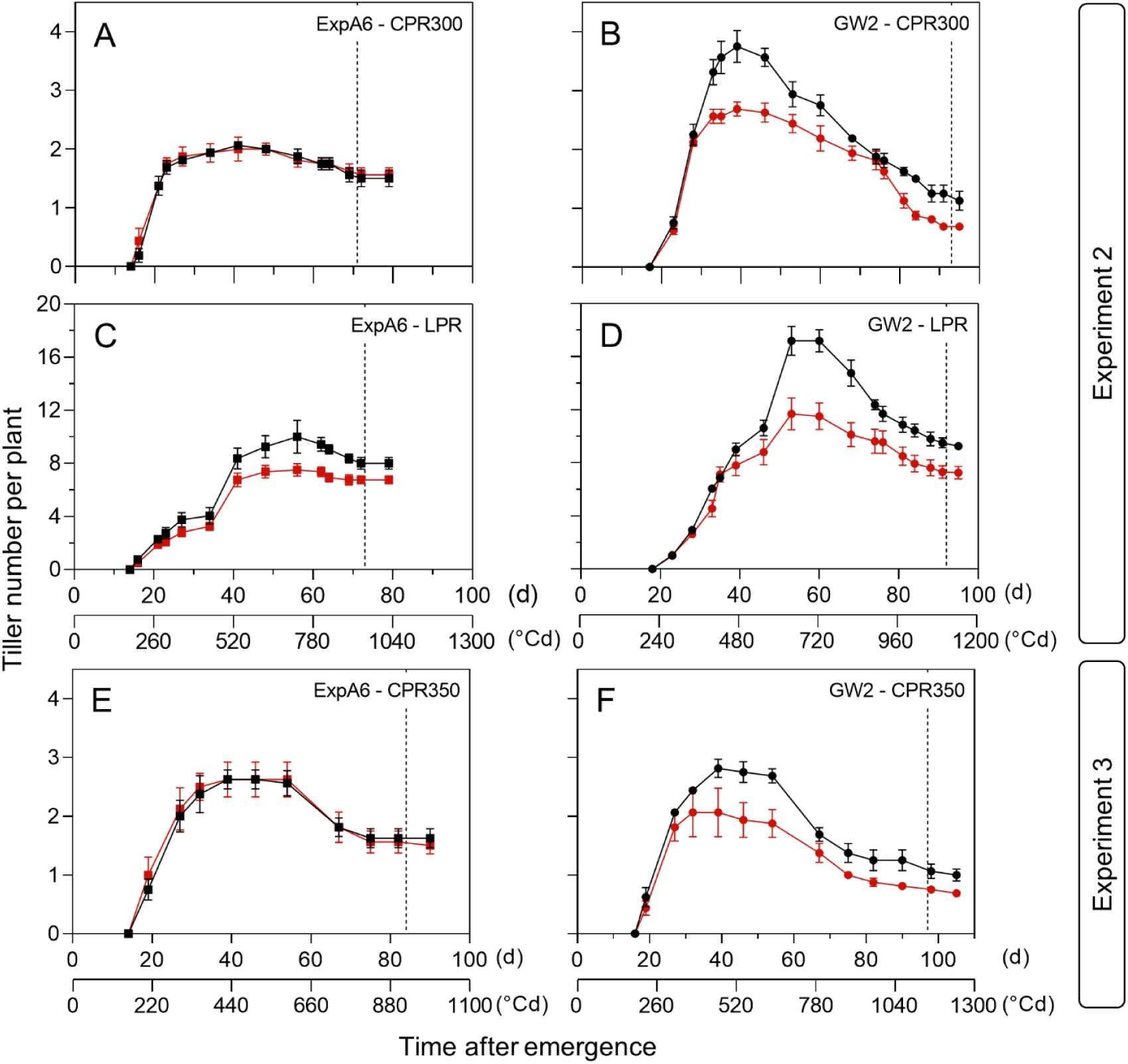
Time-course of tiller number per plant in ExpA6 lines (A, C, E) and GW2 lines (B, D, F) in experiments 2 and 3. Modified lines (i.e. *Ta*ExpA6 and *Ta*GW2) and their respective WTs are depicted by red and black symbols, respectively. In all cases, bars show the standard error of the mean. Vertical dashed line indicates the timing of anthesis.

**Table 5.**
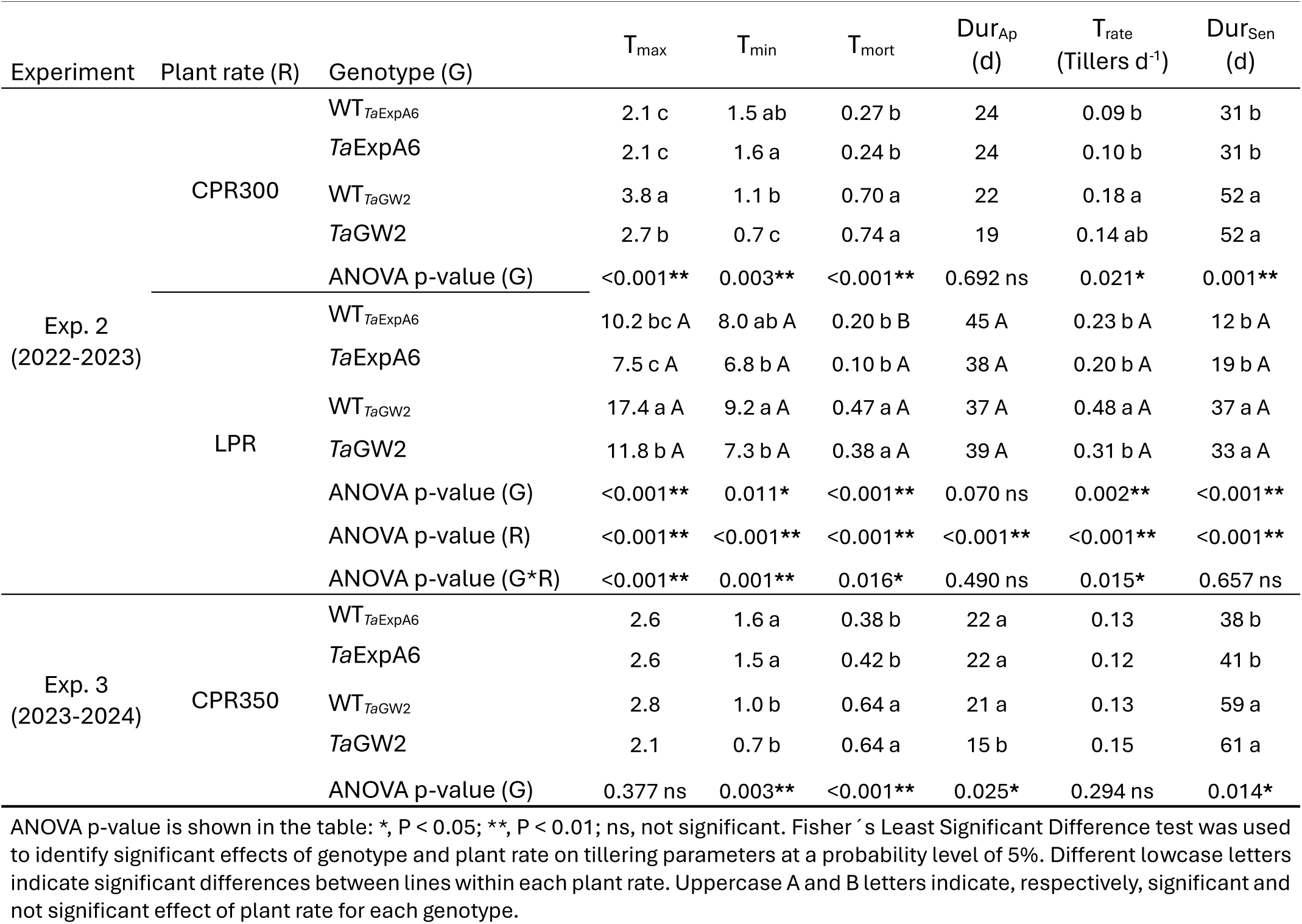
Maximum tiller number per plant (T_max_), final tiller number per plant (T_fin_), tiller mortality (T_mort_), duration of tiller appearance (Dur_Ap_) and senescence (Dur_Sen_), and tillering rate (T_rate_) recorded in ExpA6 and GW2 groups in Experiments 2 and 3.

Low plant rate treatment in Exp. 2 significantly extended Dur_Ap_ over the CPR300 treatment by 18 d (200-230 °Cd; Table 5). In addition, T_rate_ was also higher (p<0.05), whereas T_mort_ and Dur_Sen_ were lower (p< 0.05) at LPR than under CPR300 (Table 5). Consequently, NT_max_ and NT_fin_ were increased (p<0.05) 4.3-fold (11.7 vs 2.7 tillers pl^-1^) and 6.5-fold (7.8 vs 1.2 tillers pl^-1^), respectively, under LPR (Table 5).

### 3.7 Floret development underlies the differential setting of GNS observed in genotypes

In Exp. 1, floret development was similar (p>0.05) between the line *Ta*P1xGW2A and its WT at both plant rate treatments, and floret primordia fully developed up to F5 in both lines (Fig. 6B; Table S4). Correspondingly, floret development was similar between the ExpA6 lines irrespective of the experiment or treatment (Fig. 6A, C, E; Table S4). On the contrary, and supporting the relevance of this trait, the triple mutant line of the *TaGW2* gene reached lower (p<0.05) developmental rate and floret set than its WT in the most distal floret positions (i.e. F5 to F7; Fig. 6D, F). This difference was recorded across experiments and plant rates (Table S4), evidencing the remarkable trade-off between GW and GN of this line.

**Figure 6.**
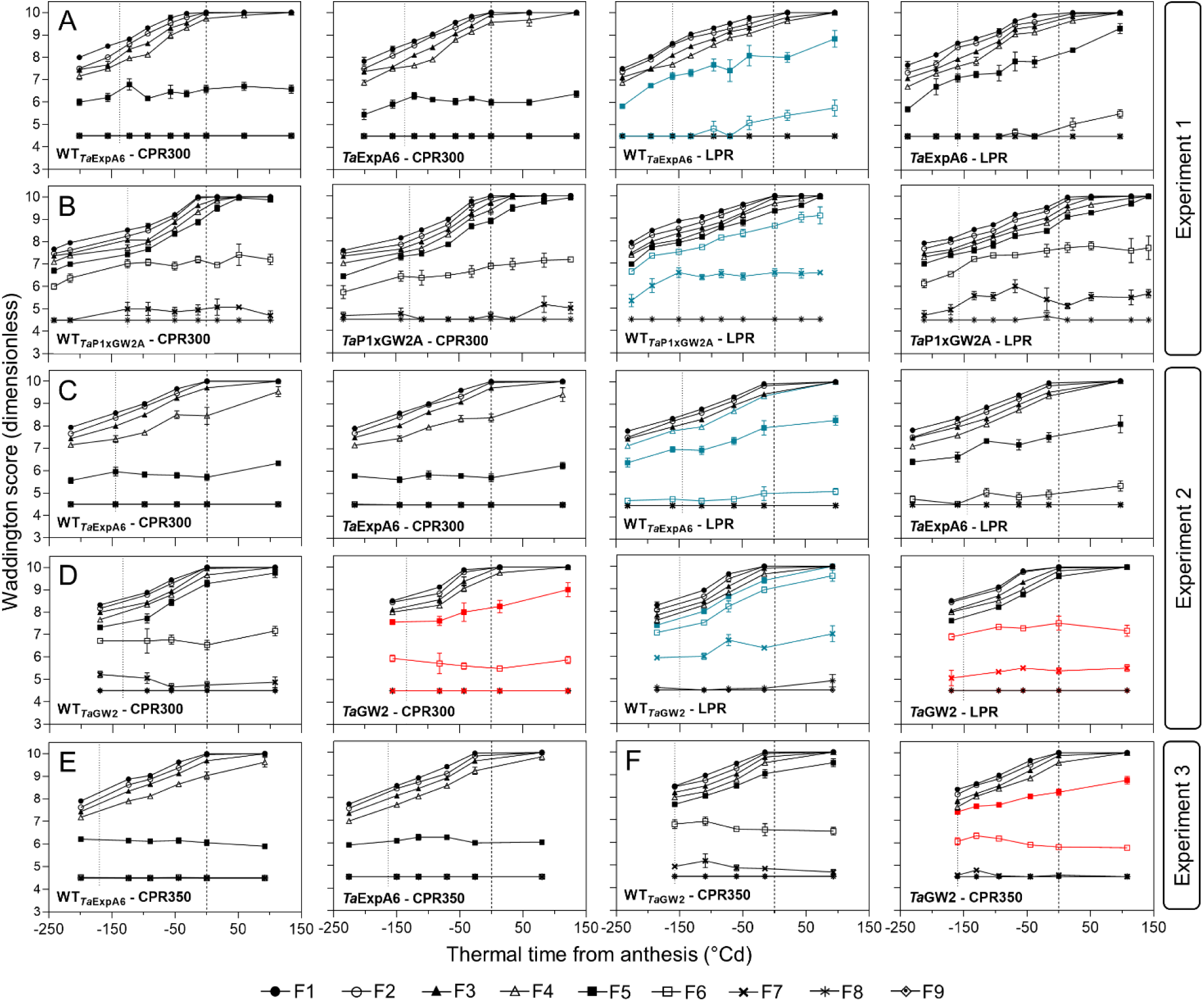
Floret development dynamics from floret 1 (F1, closest to the rachis) to floret 9 (F9, most distal to the rachis) of central spikelets of main stem spikes. (A) ExpA6 lines and (B) P1xGW2A lines in experiment 1. (C) ExpA6 lines and (D) GW2 lines in experiment 2. (E) ExpA6 lines and (F) GW2 lines in experiment 3. Bars represent the standard error of the means. Dotted and dashed vertical lines denote the timing of booting and anthesis, respectively. Floret positions significantly affected by genotype are depicted in red. Floret positions significantly affected by plant rate in each genotype group are depicted in blue in the WT figure.

The effect of LPR (44 pl m^-2^) was also evident in distal florets F5 to F7, as they always reached a higher (p<0.05) developmental stage than under the CPR treatment across lines and experiments (Fig. 6A-D; Table S4).

In Exp. 3, fertile florets per spike were quantified in main stems at anthesis (FFlo_An_), showing a close association between GNS and floret number (R^2^ = 0.98; p<0.05). Differences in FFlo_An_ were well described by the dynamics of floret development from one week before booting to 7 DAA. Remarkably, the ExpA6 lines showed similar (p>0.05) FFlo_An_ between them (Fig. 7), while this trait was lower (p<0.05) in the *Ta*GW2 triple mutant line than in its WT (i.e. 57 vs 62.8; Fig. 7), explaining the lower GNS achieved by the mutant line.

**Figure 7.**
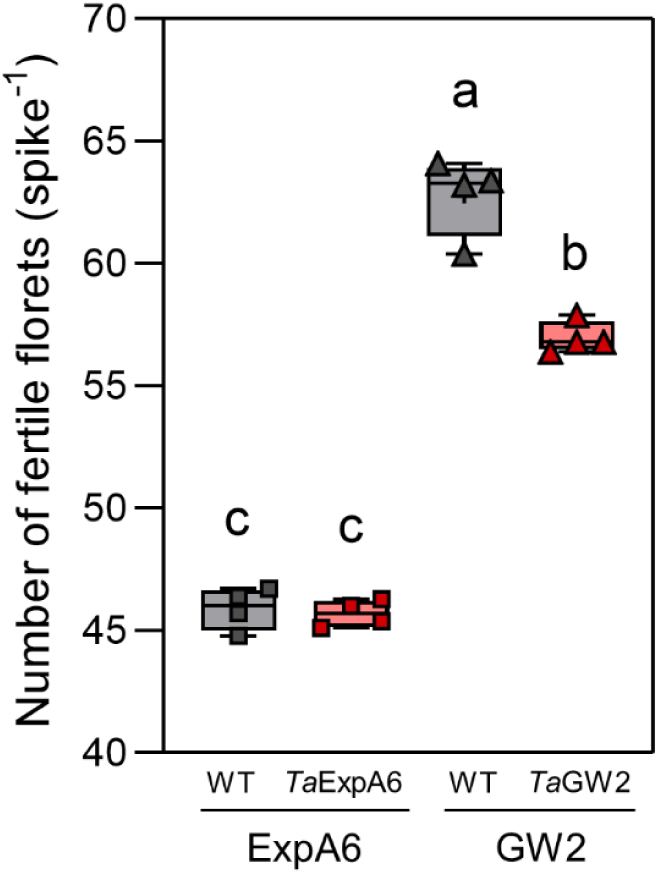
Number of fertile florets at anthesis in main stem spikes of lines *Ta*ExpA6 and *Ta*GW2 and their respective WTs in experiment 3. Whiskers denote minimum and maximum values, and median is denoted by a solid line. One-way ANOVA was performed with genotype as fixed effect. Different letters indicate significant differences between lines (p<0.05) according to Fisheŕs Least Significant Difference test post-hoc.

## 4. Discussion

The improvement of TGW through molecular and genomic tools has been highlighted as a stable goal for GY increase in wheat breeding (Jamil *et al*., 2025). Our aim in the present study was to expand the understanding of the physiological mechanisms regulating GW and the interactions between GW and GN in wheat.

As expected, lines *Ta*ExpA6 and *Ta*GW2 successfully increased TGW compared to their WTs, in agreement with previous assessments of these wheat lines (Wang *et al*., 2018; Calderini *et al*., 2021; Vicentin *et al*., 2024); while the *Ta*P1xGW2A line did not show an advantage over its WT, contrary to previous reports (Adamski *et al*., 2021).

However, grain weight improvement by *Ta*ExpA6 and *Ta*GW2 lines was only translated into a yield advantage over its WT in the *Ta*ExpA6 line under farmers plant rate, validating the effectiveness of the *TaExpA6* construct to improve GW without a trade-off with GN (Calderini *et al*., 2021; Vicentin *et al*., 2024). On the other hand, the substantial GW increase recorded in line *Ta*GW2 was counterbalanced by the reduction of both SpN and GNS across experiments, as it was also reported recently by Simmonds *et al*. (2025, Preprint) in multiple-environment field trials, including contrasting sowing-density treatments. To the best of our knowledge, the study by Simmonds *et al*. (2025, Preprint) and ours are the first reports showing a full negative compensation by GN when GW is increased through the triple mutation of the *TaGW2* gene in wheat. In rice, where the *GW2* gene was first discovered (Song *et al*., 2007), genetic manipulation of its orthologue *OsGW2* has yielded contradictory results regarding the trade-off between GN and GW. For instance, when individual plants were grown in field conditions, Huang *et al*. (2022) reported an increase in TGW and GY per plant of 19 and 9%, respectively, surprisingly without differences in GN. In contrast, other evaluations showed a clear reduction in GN in rice, which compensated for the increase in TGW (Song *et al*., 2007; Achary and Reddy, 2021).

The trade-off between both major yield components is recurrently observed in wheat (Gambín and Borrás, 2010; Golan *et al*., 2019; Xie and Sparkes, 2021; Jian *et al*., 2024; Serrago *et al*., 2025), highlighting the need to understand its underlying causes to successfully improve grain yield. This is reinforced by a recently published meta-analysis showing that the simultaneous increase of GN and GW is the most effective strategy to improve grain yield of wheat (Xi *et al*., 2024).

Notably, the triple mutation of the *TaGW2* gene increased GW in both proximal and distal grain positions of the spike (Vicentin *et al*., 2024), indicating that the observed TGW improvement found in the line *Ta*GW2 cannot be explained by a reduced number of smaller grains set in distal positions of the spike (Acreche and Slafer, 2006). The assessment of physiological determinants of GW and GN carried out in this study, allowed us to validate the physiological mechanism underpinning their negative relationship due to the overlapping of GN and GW determinations (Calderini *et al*., 2021; Vicentin *et al*., 2024), as it is pointed out below.

Regarding GW, the ovary weight at pollination (W10) has been identified as one of the main physiological drivers of GW potential in wheat (Hasan *et al*., 2011; Slafer *et al*., 2021; Zhang *et al*., 2024). The close positive association between these traits found in the present study has been widely documented in wheat (Calderini *et al*., 1999; Calderini and Reynolds, 2000; Xie *et al*., 2015; Yu *et al*., 2015; Simmonds *et al*., 2016) and further evidenced in barley (Scott *et al*., 1983; Yang *et al*., 2023), sorghum (Yang *et al*., 2009) and sunflower (Castillo *et al*., 2017). Interestingly, the ovary weight was significantly increased in the triple mutant line over its WT. Increased carpel size around anthesis by the *TaGW2* knockout mutant had already been reported (Simmonds *et al*., 2016; Vicentin *et al*., 2024). Likely, this increase is achieved through enhanced cell division and expansion in the ovary, consistent with previous findings showing that loss of *TaGW2* function increased cell number and length in the pericarp (Zhang *et al*., 2018), a maternal tissue derived from the ovary.

The assessment of ovary growth carried out in our study expands this knowledge by showing that the effect of *TaGW2* loss of function on ovary weight is already significant at booting, and then maintained during the Bo-W10 period through an increased growth rate. This early impact of *TaGW2* disruption on ovary growth is in agreement with the gene expression profile found in young developing spikes (Ramírez-González *et al*., 2018; Tillett *et al*., 2022 and references therein), and explains the high impact on GW reported for this gene knockout in our and previous studies (Wang *et al*., 2018; Vicentin *et al*., 2024). On the contrary, no effect of the *TaExpA6* construct was observed on ovary weight from booting to pollination respect to its WT. In fact, the positive impact of the *TaExpA6* construct on GW was detected by two weeks after anthesis (Vicentin *et al*., 2024). Therefore, both the positive effect *TaGW2* disruption on ovary size and the lack of effect by the *TaExpA6* construct validate the proof of concept reported by Calderini *et al*. (2021) proposing the overlapping between grain number and grain weight determinations as the cause of the trade-off between both main yield components.

About GN, the development of florets during the window from Bo to W10, especially in distal positions of the spikelet, is a key for GN determination (Slafer and Andrade, 1993; Miralles *et al*., 1998; González *et al*., 2005; González-Navarro *et al*., 2015; Ferrante *et al*., 2017; Ferrante *et al*., 2020), and it has been associated to the assimilates allocated to the spike (Fischer, 1985; Slafer and Andrade, 1993; Abbate *et al*., 1995; González *et al*., 2005; Ferrante *et al*., 2010; 2013; Zhu *et al*., 2022; Bicego *et al*., 2024). Interestingly, a lower developmental rate of distal florets (i.e. positions 5 to 7) was observed in the present study under the effect of the *TaGW2* disruption. Thus, we hypothesize that the increase in ovary weight in proximal floret positions induced by the mutation of *TaGW2* gene might reduce the allocation of resources to distal, more labile, floret primordia, decreasing the likelihood of reaching W10 at pollination, and the number of fertile florets at anthesis (FFlo_An_). As a consequence, GNS is reduced, as it was recorded in the *Ta*GW2 line in the present study. This hypothesis is in agreement with the fate of distal florets reported by Guo *et al*. (2016). However, the weight of ovaries from positions F4 and F5 at pollination was also increased in the triple mutant line when compared to its WT (Vicentin *et al*., 2024). This discrepancy most likely arises from the sampling procedure, which could bias the results given that only ovaries that reached the W10 stage were measured. As these ovaries were a low proportion of the total, they did not accurately reflect the average ovary weight at these positions.

It has been shown that floret survival and GN are positively associated with the duration of the spike growth period (i.e. from terminal spikelet to anthesis; Ds), by increasing the likelihood of distal florets to become fertile (Miralles *et al*., 2000; González-Navarro *et al*., 2015; Fischer *et al*., 2024). Empirical evidences support that it leads to enhanced GNS (Wang *et al*., 2017; Ochagavía *et al*., 2018; Glenn *et al*., 2022; Slafer *et al*., 2022a; Kronenberg *et al*., 2025, Preprint). Therefore, the negative effect of the *TaGW2* triple mutant line on GNS could be ameliorated by extending Ds.

In addition to GNS, the disruption of *TaGW2* also decreased SpN compared to its WT in our study. This negative impact is likely the result of a pleiotropic effect of *TaGW2* gene knock-out. This gene codes for a RING-type E3 ligase, which mediates proteolysis through the ubiquitin-proteasome pathway (Song *et al*., 2007), and is constitutively expressed across wheat tissues from emergence to maturity, including apical meristems during tillering (Ramírez-González *et al*., 2018; Fig. S5). Recently, Jian *et al*. (2024) showed that the *TaGW2* gene positively regulates tillering by mediating the degradation of *TaSPL14*, a negative regulator of tillering in wheat and rice (Jiao *et al*., 2010; Cao *et al*., 2023). Remarkably, lower number of spikes per plant was also reported in barley lines with targeted mutations of the *GW2* gene (Kis *et al*., 2024). At the best of our knowledge, there are not studies reporting the effect of the *OsGW2* gene knock out on tillering of rice, reinforcing the importance of the tillers time-course showed in the present study (Fig. 6).

The negative effect of *TaGW2* disruption on both SpN and GNS emphasize the relevance of the specificity of the *PinB* promoter of the *TaExpA6* construct driving the expression of the gene in grain tissues (Gautier *et al*., 1994; Digeon *et al*., 1999), and its post-anthesis timing avoiding the overlapping between GW and GN determinations, dodging the trade-offs as a consequence (Calderini *et al*., 2021).

Supporting this, *TaCYP78A5* overexpression led to increased GW and yield only when it was restricted to grain maternal tissues, whereas the constitutive overexpression of this gene had a detrimental effect on tillering, offsetting yield gains (Guo *et al*., 2022).

Contrasting plant rate treatments carried out in this study and by Simmonds *et al*. (2025, Preprint) provide valuable insights into the response of SpN, GNS and GW and their trade-offs. In our study, the significant GxE interaction was observed for GY response to the 8-fold reduction in plant number, was mainly explained by the differential tillering potential among genotypes, as in previous studies (Frederick and Marshall, 1985; Valério *et al*., 2009; Lollato *et al*., 2024). The positive effect of the *TaExpA6* construct on GY was neutralized under the LPR treatment, due to reductions in SpN. This is a likely consequence of the shortened Em-An period of Fielder background (Fig. 1), and not due to a negative impact of the construct *per se*, regarding the lower SpN and GY were also observed under LPR for the WT*_Ta_*_ExpA6_ line. On the other hand, the greater reliance of GN on tiller spikes under LPR amplified the negative effect of *TaGW2* mutation on SpN. Although the increase of plant rate, to minimize the contribution of tillers to total GN and yield, could be a promising strategy to counteract the negative effect the *TaGW2* knock-out on SpN, Simmonds *et al*. (2025, Preprint) demonstrated its lack of effectiveness by trying 500 pl m^-2^, which failed to achieve any yield advantage over the WT.

LPR led to a simultaneous increase of GW and GNS in main stems and tillers compared to the CPR, consistent with previous evaluations of winter and spring wheat cultivars (Li *et al*., 2016; Hasan *et al*., 2022). Thus, an uneven trade-off between GW and GNS was observed when GW was improved genetically (*TaGW2*) or by management (LPR). As observed in line *Ta*GW2, the increase of individual GW in LPR was associated with a positive effect of this treatment on the ovary weight at pollination, which was achieved by both a higher growth rate and duration of the Bo-W10 period. However, despite the close association between final GW and ovary weight at W10 at individual positions, overall average GW did not show differences between LPR and CPR. The lack of impact of individual GW improvement on TGW under LPR can be attributed to (i) the higher GNS resulting in an increased proportion of grains from distal positions of the spike (Acreche and Slafer, 2006), and (ii) the increased proportion of grains from tillers, both of which have an intrinsic lower GW.

Unlike the *TaGW2* mutation, the LPR treatment also had a positive effect on floret development in distal positions of the spikelet. Former studies, such as Bustos *et al*. (2013) and Bicego *et al*. (2024), showed a positive response of the rate of floret survival when the resource availability per spike were increased by reducing the plant rate. These results support that a greater resource allocation to the spike would increase the likelihood of improving both ovary growth and floret fertility, mitigating the trade-off between GW and GNS observed in line *Ta*GW2. Slafer *et al*. (2022b) explained the relationship among GN subcomponents by the same mechanism.

Nevertheless, the lower floret survival in the *Ta*GW2 line occurred even under increased resource availability to the spike at LPR. This agrees with Jones *et al*. (2021), who showed that the response to allelic variants of *TaGW2-A* is not modified by the availability of resources. Most likely, the intra-spike partitioning of resources in favour of structural components of the spike could contribute to the reduced FE observed in the *Ta*GW2 triple mutant line, in addition to the increased ovary size, as inferred from the findings of Slafer *et al*. (2015).

### 4.1 Concluding remarks

The present study confirmed the effectiveness of the *TaExpA6* construct to increase GW without a trade-off with GN, as opposed to the triple knock-out of the *TaGW2* gene, which markedly increased GW but was counteracted by the fall of both SpN and GNS, resulting in null yield advantage. These contrasting outcomes might stem from differences between these gene’s expression patterns and physiological roles: i.e., the *TaExpA6* construct acts in post-anthesis in grain tissues, avoiding the overlap with GN determination, while *TaGW2* loss-of-function (i) restricts tiller generation likely due to a pleiotropic effect and (ii) promotes early ovary growth at the expense of distal floret development, ultimately decreasing GNS. The increased resource allocation to the spikes under a low plant rate of 44 pl m^-2^ improved both ovary growth and distal floret development across genotypes, alleviating the GW–GNS trade-off. However, this strategy did not modify the negative impact of *TaGW2* mutation, indicating a genetic limitation to overcome this trade-off. On the other hand, the positive effect of *TaExpA6* overexpression on GY was plant-rate-dependent, likely due to the low tillering potential of the Fielder background, which constrained SpN under LPR. Altogether, these findings underscore the importance of considering the interactions among traits for GY improvement in wheat, and the relevance of the timing of gene expression and its tissue specificity to avoid the trade-off between GW and GN when designing molecular strategies aimed at increasing GW.

## 6. Acknowledgements

The *Ta*ExpA6 line and its segregant WT were developed by Dr. Emma Wallington from the National Institute of Agricultural Botany (NIAB), UK (Calderini *et al*., 2021). Japan Tobacco Inc. is recognized as a holder for the technology included in PureWheat® licensed from Japan Tobacco Inc. *Ta*GW2 and *Ta*P1xGW2A lines were generously provided by Prof. Cristóbal Uauy from John Innes Centre (UK). We also thank the experimental field staff of the Austral Farming Experimental Station of Universidad Austral de Chile for the technical assistance provided. Technical contributions and management of field trials by Marcelo Castro Moraga and Cristóbal Castro Villalón is truly appreciated. We are also very grateful to María Beatriz Ugalde Jaramillo and Dr. Anita Arenas Miranda (Plant Nutrition and Genomics Lab – UACh) for performing the molecular analysis included in this study. The work on samples processing by Beatriz Shibar (UACh) and Magda Lobnik is recognized.

## 7. Author contributions

**DFC** conceived the study, coordinated experiments and data analysis; **LV** conducted field experiments and formal analysis of the data. **LV** and **DFC** wrote the manuscript.

## 10. Data availability

All data supporting the findings of this study will be made available on request.

## References

Abbate, P.E., Andrade, F.H., and Culot, J.P. 1995. The effects of radiation and nitrogen on number of grains in wheat. The Journal of Agricultural Science 124, 351–360.

Achary, V.M.M., and Reddy, M.K. 2021. CRISPR-Cas9 mediated mutation in GRAIN WIDTH and WEIGHT2 (GW2) locus improves aleurone layer and grain nutritional quality in rice. Sci Rep 11, 21941.

Acreche, M.M., and Slafer, G.A. 2006. Grain weight response to increases in number of grains in wheat in a Mediterranean area. Field Crops Research 98, 52–59.

Adamski, N.M., Simmonds, J., Brinton, J.F., et al. 2021. Ectopic expression of Triticum polonicum VRT-A2 underlies elongated glumes and grains in hexaploid wheat in a dosage-dependent manner. Plant Cell 33, 2296–2319.

Bicego, B., Savin, R., Girousse, C., Allard, V., and Slafer, G.A. 2024. Tillering and floret development dynamics in wheat cultivars of contrasting spike fertility plasticity. Field Crops Research 319, 109654.

Borrás, L., Slafer, G.A., and Otegui, M.a.E. 2004. Seed dry weight response to source–sink manipulations in wheat, maize and soybean: a quantitative reappraisal. Field Crops Research 86, 131–146.

Borrill, P., Fahy, B., Smith, A.M., and Uauy, C. 2015. Wheat Grain Filling Is Limited by Grain Filling Capacity rather than the Duration of Flag Leaf Photosynthesis: A Case Study Using NAM RNAi Plants. PLoS One 10, e0134947.

Braun, H., Atlin, G., and Payne, T. 2010. Multi-location testing as a tool to identify plant response to global climate change. CABI International.

Brinton, J., Simmonds, J., Minter, F., Leverington-Waite, M., Snape, J., and Uauy, C. 2017. Increased pericarp cell length underlies a major quantitative trait locus for grain weight in hexaploid wheat. New Phytol 215, 1026–1038.

Brunner, S., Weichert, H., Meissle, M., Romeis, J., and Weber, H. 2024. Field trials reveal trade-offs between grain size and grain number in wheat ectopically expressing a barley sucrose transporter. Field Crops Research 316, 109506.

Bustos, D.V., Hasan, A.K., Reynolds, M.P., and Calderini, D.F. 2013. Combining high grain number and weight through a DH-population to improve grain yield potential of wheat in high-yielding environments. Field Crops Research 145, 106–115.

Calderini, D.F., Abeledo, L.G., Savin, R., and Slafer, G.A. 1999. Effect of temperature and carpel size during pre-anthesis on potential grain weight in wheat. The Journal of Agricultural Science 132, 453–459.

Calderini, D.F., Castillo, F.M., Arenas, M.A., et al. 2021. Overcoming the trade-off between grain weight and number in wheat by the ectopic expression of expansin in developing seeds leads to increased yield potential. New Phytol 230, 629–640.

Calderini, D.F., Dreccer, M.F., and Slafer, G.A. 1995. Genetic improvement in wheat yield and associated traits. A re-examination of previous results and the latest trends. Plant Breeding 114, 108–112.

Calderini, D.F., and Reynolds, M.P. 2000. Changes in grain weight as a consequence of de-graining treatments at pre- and post-anthesis in synthetic hexaploid lines of wheat (Triticum durum x T. tauschii). Functional Plant Biology 27.

Calderini, D.F., Reynolds, M.P., and Slafer, G.A. 2006. Sourcesink effects on grain weight of bread wheat, durum wheat, and triticale at different locations. Australian Journal of Agricultural Research 57, 227–233.

Cao, L., Li, T., Geng, S., Zhang, Y., Pan, Y., Zhang, X., Wang, F., and Hao, C. 2023. TaSPL14-7A is a conserved regulator controlling plant architecture and yield traits in common wheat (Triticum aestivum L.). Frontiers in Plant Science Volume 14–2023.

Castillo, F.M., Vásquez, S.C., and Calderini, D.F. 2017. Does the pre-flowering period determine the potential grain weight of sunflower? Field Crops Research 212, 23–33.

Di Rienzo, J.A., Casanoves, F., Balzarini, M.G., Gonzalez, L., Tablada, M., and Robledo, C.W. 2020. InfoStat [Online]. Córdoba, Argentina: Universidad Nacional de Córdoba. Available: http://www.infostat.com.ar/ [Accessed 11 March 2023].

Digeon, J.-F., Guiderdoni, E., Alary, R., Michaux-Ferrière, N., Joudrier, P., and Gautier, M.-F. 1999. Cloning of a wheat puroindoline gene promoter by IPCR and analysis of promoter regions required for tissue-specific expression in transgenic rice seeds. Plant Molecular Biology 39, 1101–1112.

Dreccer, M.F., Grashoff, C., and Rabbinge, R. 1997. Source-sink ratio in barley (Hordeum vulgare L.) during grain filling: effects on senescence and grain protein concentration. Field Crops Research 49, 269–277.

Dreccer, M.F., Zwart, A.B., Schmidt, R.C., Condon, A.G., Awasi, M.A., Grant, T.J., Galle, A., Bourot, S., and Frohberg, C. 2022. Wheat yield potential can be maximized by increasing red to far-red light conditions at critical developmental stages. Plant Cell Environ 45, 2652–2670.

Ellen, J. 1990. Effects of Nitrogen and Plant Density on Growth, Yield and Chemical Composition of Two Winter Wheat (Triticum aestivum L.) Cultivars. Journal of Agronomy and Crop Science 164, 174–183.

Evers, J.B., Vos, J., Andrieu, B., and Struik, P.C. 2006. Cessation of tillering in spring wheat in relation to light interception and red : far-red ratio. Ann Bot 97, 649–658.

Fao. 2017. The future of food and agriculture - Trends and challenges. Rome.

Ferrante, A., Cartelle, J., Savin, R., and Slafer, G.A. 2017. Yield determination, interplay between major components and yield stability in a traditional and a contemporary wheat across a wide range of environments. Field Crops Research 203, 114–127.

Ferrante, A., Savin, R., and Slafer, G.A. 2010. Floret development of durum wheat in response to nitrogen availability. J Exp Bot 61, 4351–4359.

Ferrante, A., Savin, R., and Slafer, G.A. 2013. Floret development and grain setting differences between modern durum wheats under contrasting nitrogen availability. J Exp Bot 64, 169–184.

Ferrante, A., Savin, R., and Slafer, G.A. 2020. Floret development and spike fertility in wheat: Differences between cultivars of contrasting yield potential and their sensitivity to photoperiod and soil N. Field Crops Research 256.

Fischer, R.A. 1985. Number of kernels in wheat crops and the influence of solar radiation and temperature. The Journal of Agricultural Science 105, 447–461.

Fischer, R.A., Byerlee, D., and Edmeades, G.O. 2014. Crop Yields and Global Food Security - Will yield increase continue to feed the world? ACIAR Monograph No. 158. Canberra: Australian Centre for International Agricutural Research.

Fischer, T., Gonzalez, F.G., and Miralles, D.J. 2024. Breeding for increased grains/m2 in wheat crops through targeting critical period duration: A review. Field Crops Research 316, 109497.

Frederick, J.R., and Marshall, H.G. 1985. Grain Yield and Yield Components of Soft Red Winter Wheat as Affected by Management Practices1. Agronomy Journal 77, 495–499.

Gambín, B.L., and Borrás, L. 2010. Resource distribution and the trade-off between seed number and seed weight: a comparison across crop species. Annals of Applied Biology 156, 91–102.

Gautier, M.-F., Aleman, M.-E., Guirao, A., Marion, D., and Joudrier, P. 1994. Triticum aestivum puroindolines, two basic cystine-rich seed proteins: cDNA sequence analysis and developmental gene expression. Plant Molecular Biology 25, 43–57.

Glenn, P., Zhang, J., Brown-Guedira, G., Dewitt, N., Cook, J.P., Li, K., Akhunov, E., and Dubcovsky, J. 2022. Identification and characterization of a natural polymorphism in FT-A2 associated with increased number of grains per spike in wheat. Theor Appl Genet 135, 679–692.

Golan, G., Ayalon, I., Perry, A., Zimran, G., Ade-Ajayi, T., Mosquna, A., Distelfeld, A., and Peleg, Z. 2019. GNI-A1 mediates trade-off between grain number and grain weight in tetraploid wheat. Theor Appl Genet 132, 2353–2365.

González-Navarro, O.E., Griffiths, S., Molero, G., Reynolds, M.P., and Slafer, G.A. 2015. Dynamics of floret development determining differences in spike fertility in an elite population of wheat. Field Crops Research 172, 21–31.

González, F.G., Slafer, G.A., and Miralles, D.J. 2003. Floret development and spike growth as affected by photoperiod during stem elongation in wheat. Field Crops Research 81, 29–38.

González, F.G., Slafer, G.A., and Miralles, D.J. 2005. Floret development and survival in wheat plants exposed to contrasting photoperiod and radiation environments during stem elongation. Functional Plant Biology 32, 189–197.

Guo, L., Ma, M., Wu, L., et al. 2022. Modified expression of TaCYP78A5 enhances grain weight with yield potential by accumulating auxin in wheat (Triticum aestivum L.). Plant Biotechnol J 20, 168–182.

Guo, Z., Slafer, G.A., and Schnurbusch, T. 2016. Genotypic variation in spike fertility traits and ovary size as determinants of floret and grain survival rate in wheat. Journal of Experimental Botany 67, 4221–4230.

Hasan, A.K., Carrasco-G, F.E., Lizana, C.X., and Calderini, D.F. 2022. Low seed rate in square planting arrangement has neutral or positive effect on grain yield and improves grain nitrogen and phosphorus uptake in wheat. Field Crops Research 288.

Hasan, A.K., Herrera, J., Lizana, C., and Calderini, D.F. 2011. Carpel weight, grain length and stabilized grain water content are physiological drivers of grain weight determination of wheat. Field Crops Research 123, 241–247.

Huang, J., Chen, Z., Lin, J., et al. 2022. gw2.1, a new allele of GW2, improves grain weight and grain yield in rice. Plant Science 325, 111495.

Jablonski, B., Szala, K., Przyborowski, M., Bajguz, A., Chmur, M., Gasparis, S., Orczyk, W., and Nadolska-Orczyk, A. 2021. TaCKX2.2 Genes Coordinate Expression of Other TaCKX Family Members, Regulate Phytohormone Content and Yield-Related Traits of Wheat. Int J Mol Sci 22.

Jamil, M., Ahmad, W., Sanwal, M., and Maqsood, M.F. 2025. Gene editing and GWAS for digital imaging analysis of wheat grain weight, size and shape are inevitable to enhance the yield. Cereal Research Communications.

Jian, C., Pan, Y., Liu, S., et al. 2024. The TaGW2-TaSPL14 module regulates the trade-off between tiller number and grain weight in wheat. Journal of Integrative Plant Biology 66, 1953–1965.

Jiao, Y., Wang, Y., Xue, D., et al. 2010. Regulation of OsSPL14 by OsmiR156 defines ideal plant architecture in rice. Nature Genetics 42, 541–544.

Jones, B.H., Blake, N.K., Heo, H.Y., Martin, J.M., Torrion, J.A., and Talbert, L.E. 2021. Allelic response of yield component traits to resource availability in spring wheat. Theor Appl Genet 134, 603–620.

Kis, A., Polgári, D., Dalmadi, Á., Ahmad, I., Rakszegi, M., Sági, L., Csorba, T., and Havelda, Z. 2024. Targeted mutations in the GW2.1 gene modulate grain traits and induce yield loss in barley. Plant Science 340, 111968.

Kronenberg, L., Gonzalez-Navarro, O.E., Collier, S., Chhetry, M., Tailby, P., Leverington-Waite, M., Wingen, L.U., and Griffiths, S. 2025. Buying time – increasing yield potential in wheat by extending stem elongation duration. BioRxivdoi: 10.1101/2025.04.28.650957. [Preprint]

Kumar, A., Mantovani, E.E., Seetan, R., et al. 2016. Dissection of Genetic Factors underlying Wheat Kernel Shape and Size in an Elite × Nonadapted Cross using a High Density SNP Linkage Map. The Plant Genome 9, plantgenome2015.2009.0081.

Li, Y., Cui, Z., Ni, Y., Zheng, M., Yang, D., Jin, M., Chen, J., Wang, Z., and Yin, Y. 2016. Plant Density Effect on Grain Number and Weight of Two Winter Wheat Cultivars at Different Spikelet and Grain Positions. PLoS One 11, e0155351.

Lichthardt, C., Chen, T.W., Stahl, A., and Stutzel, H. 2020. Co-Evolution of Sink and Source in the Recent Breeding History of Winter Wheat in Germany. Front Plant Sci 10, 1771.

Liu, J., Chen, Z., Wang, Z., et al. 2021. Ectopic expression of VRT-A2 underlies the origin of Triticum polonicum and Triticum petropavlovskyi with long outer glumes and grains. Mol Plant 14, 1472–1488.

Lloveras, J., Manent, J., Viudas, J., López, A., and Santiveri, P. 2004. Seeding Rate Influence on Yield and Yield Components of Irrigated Winter Wheat in a Mediterranean Climate. Agronomy Journal 96, 1258–1265.

Lollato, R.P., Pradella, L.O., Giordano, N., Ryan, L.P., Soler, J.R., Simão, L.M., Jaenisch, B.R., and Horton, R. 2024. Winter wheat response to plant density in yield contest fields. Crop Science n/a.

Mcdonald, P., and Henderson, A.R. 1964. Determination of water-soluble carbohydrates in grass. Journal of the Science of Food and Agriculture 15, 395–398.

Milner, M.J., Bowden, S., Craze, M., and Wallington, E.J. 2021. Ectopic expression of TaBG1 increases seed size and alters nutritional characteristics of the grain in wheat but does not lead to increased yields. BMC Plant Biol 21, 524.

Miralles, D.J., Katz, S.D., Colloca, A., and Slafer, G.A. 1998. Floret development in near isogenic wheat lines differing in plant height. Field Crops Research 59, 21–30.

Miralles, D.J., Richards, R.A., and Slafer, G.A. 2000. Duration of the stem elongation period influences the number of fertile florets in wheat and barley. Functional Plant Biology 27, 931–940.

Molero, G., Joynson, R., Pinera-Chavez, F.J., Gardiner, L.J., Rivera-Amado, C., Hall, A., and Reynolds, M.P. 2019. Elucidating the genetic basis of biomass accumulation and radiation use efficiency in spring wheat and its role in yield potential. Plant Biotechnol J 17, 1276–1288.

Mora-Ramirez, I., Weichert, H., Von Wiren, N., Frohberg, C., De Bodt, S., Schmidt, R.C., and Weber, H. 2021. The da1 mutation in wheat increases grain size under ambient and elevated CO(2) but not grain yield due to trade-off between grain size and grain number. Plant Environ Interact 2, 61–73.

Ochagavía, H., Prieto, P., Savin, R., Griffiths, S., and Slafer, G.A. 2018. Earliness per se effects on developmental traits in hexaploid wheat grown under field conditions. European Journal of Agronomy 99, 214–223.

Okamoto, Y., and Takumi, S. 2013. Pleiotropic effects of the elongated glume gene P1 on grain and spikelet shape-related traits in tetraploid wheat. Euphytica 194, 207–218.

Qin, L., Hao, C., Hou, J., Wang, Y., Li, T., Wang, L., Ma, Z., and Zhang, X. 2014. Homologous haplotypes, expression, genetic effects and geographic distribution of the wheat yield gene TaGW2. BMC Plant Biol 14, 107.

Quintero, A., Molero, G., Reynolds, M.P., and Calderini, D.F. 2018. Trade-off between grain weight and grain number in wheat depends on GxE interaction: A case study of an elite CIMMYT panel (CIMCOG). European Journal of Agronomy 92, 17–29.

Ramírez-González, R.H., Borrill, P., Lang, D., et al. 2018. The transcriptional landscape of polyploid wheat. Science 361, eaar6089.

Ray, D.K., Mueller, N.D., West, P.C., and Foley, J.A. 2013. Yield Trends Are Insufficient to Double Global Crop Production by 2050. PLoS One 8, e66428.

Reynolds, M.P., Foulkes, M.J., Slafer, G.A., Berry, P., Parry, M.a.J., Snape, J.W., and Angus, W.J. 2009. Raising yield potential in wheat. Journal of Experimental Botany 60, 1899–1918.

Richards, R.A. (1996). Increasing yield potential in wheat—source and sink limitations. In Increasing Yield Potential in Wheat: Breaking the Barriers, eds. M.P. Reynolds, S. Rajaram & A. Mcnab. (Mexico City: CIMMYT), 134–149.

Rivera-Amado, C., Trujillo-Negrellos, E., Molero, G., Reynolds, M.P., Sylvester-Bradley, R., and Foulkes, M.J. 2019. Optimizing dry-matter partitioning for increased spike growth, grain number and harvest index in spring wheat. Field Crops Research 240, 154–167.

Scott, W.R., Appleyard, M., Fellowes, G., and Kirby, E.J.M. 1983. Effect of genotype and position in the ear on carpel and grain growth and mature grain weight of spring barley. The Journal of Agricultural Science 100, 383–391.

Serrago, R.A., Alzueta, I., Savin, R., and Slafer, G.A. 2013. Understanding grain yield responses to source–sink ratios during grain filling in wheat and barley under contrasting environments. Field Crops Research 150, 42–51.

Serrago, R.A., García, G.A., Savin, R., Miralles, D.J., and Slafer, G.A. 2025. Relevance of grain number and grain weight on barley yield responses to environmental and genetic factors. Field Crops Research 328, 109922.

Sestili, F., Pagliarello, R., Zega, A., et al. 2019. Enhancing grain size in durum wheat using RNAi to knockdown GW2 genes. Theor Appl Genet 132, 419–429.

Simmonds, J., Crane, P., Eade, S., et al. 2025. Increased grain weight conferred by GW2 mutations in wheat does not translate into yield gains in multi-year field trials of near-isogenic lines. BioRxiv doi: [Preprint]

Simmonds, J., Scott, P., Brinton, J., Mestre, T.C., Bush, M., Del Blanco, A., Dubcovsky, J., and Uauy, C. 2016. A splice acceptor site mutation in TaGW2-A1 increases thousand grain weight in tetraploid and hexaploid wheat through wider and longer grains. Theor Appl Genet 129, 1099–1112.

Slafer, G.A., and Andrade, F.H. 1993. Physiological attributes related to the generation of grain yield in bread wheat cultivars released at different eras. Field Crops Research 31, 351–367.

Slafer, G.A., Elia, M., Savin, R., García, G.A., Terrile, I.I., Ferrante, A., Miralles, D.J., and González, F.G. 2015. Fruiting efficiency: an alternative trait to further rise wheat yield. Food and Energy Security 4, 92–109.

Slafer, G.A., Foulkes, M.J., Reynolds, M.P., Murchie, E.H., Carmo-Silva, E., Flavell, R., Gwyn, J., Sawkins, M., and Griffiths, S. 2022a. A ‘wiring diagram’ for sink strength traits impacting wheat yield potential. Journal of Experimental Botany 74, 40–71.

Slafer, G.A., Foulkes, M.J., Reynolds, M.P., Murchie, E.H., Carmo-Silva, E., Flavell, R., Gwyn, J., Sawkins, M., and Griffiths, S. 2023. A ‘wiring diagram’ for sink strength traits impacting wheat yield potential. Journal of Experimental Botany 74, 40–71.

Slafer, G.A., García, G.A., Serrago, R.A., and Miralles, D.J. 2022b. Physiological drivers of responses of grains per m2 to environmental and genetic factors in wheat. Field Crops Research 285, 108593.

Slafer, G.A., and Savin, R. 1994. Source—sink relationships and grain mass at different positions within the spike in wheat. Field Crops Research 37, 39–49.

Slafer, G.A., Savin, R., Pinochet, D., and Calderini, D.F. (2021). Wheat. In Crop Physiology. Case Histories for Major Crops, eds. V. Sadras & D.F. Calderini. (Elsevier, London: Academic Press), 99-145.

Slafer, G.A., Savin, R., and Sadras, V.O. 2014. Coarse and fine regulation of wheat yield components in response to genotype and environment. Field Crops Research 157, 71–83.

Song, X.J., Huang, W., Shi, M., Zhu, M.Z., and Lin, H.X. 2007. A QTL for rice grain width and weight encodes a previously unknown RING-type E3 ubiquitin ligase. Nat Genet 39, 623–630.

Su, Z., Hao, C., Wang, L., Dong, Y., and Zhang, X. 2011. Identification and development of a functional marker of TaGW2 associated with grain weight in bread wheat (Triticum aestivum L.). Theor Appl Genet 122, 211–223.

Tillett, B.J., Hale, C.O., Martin, J.M., and Giroux, M.J. 2022. Genes Impacting Grain Weight and Number in Wheat (Triticum aestivum L. ssp. aestivum). Plants (Basel) 11.

Ugarte, C.C., Trupkin, S.A., Ghiglione, H., Slafer, G., and Casal, J.J. 2010. Low red/far-red ratios delay spike and stem growth in wheat. J Exp Bot 61, 3151–3162.

Valério, I.P., Carvalho, F.I.D., Oliveira, A.D., et al. 2009. Seeding density in wheat genotypes as a function of tillering potential. Scientia Agricola 66.

Van Dijk, M., Morley, T., Rau, M.L., and Saghai, Y. 2021. A meta-analysis of projected global food demand and population at risk of hunger for the period 2010-2050. Nat Food 2, 494–501.

Vicentin, L., Canales, J., and Calderini, D.F. 2024. The trade-off between grain weight and grain number in wheat is explained by the overlapping of the key phases determining these major yield components. Frontiers in Plant Science Volume 15–2024.

Vollset, S.E., Goren, E., Yuan, C.W., et al. 2020. Fertility, mortality, migration, and population scenarios for 195 countries and territories from 2017 to 2100: a forecasting analysis for the Global Burden of Disease Study. Lancet 396, 1285–1306.

Waddington, S.R., Cartwright, P.M., and Wall, P.C. 1983. A Quantitative Scale of Spike Initial and Pistil Development in Barley and Wheat. Annals of Botany 51, 119–130.

Wang, W., Simmonds, J., Pan, Q., et al. 2018. Gene editing and mutagenesis reveal inter-cultivar differences and additivity in the contribution of TaGW2 homoeologues to grain size and weight in wheat. Theor Appl Genet 131, 2463–2475.

Wang, X., Dong, L., Hu, J., et al. 2019. Dissecting genetic loci affecting grain morphological traits to improve grain weight via nested association mapping. Theor Appl Genet 132, 3115–3128.

Wang, Y., Yu, H., Tian, C., Sajjad, M., Gao, C., Tong, Y., Wang, X., and Jiao, Y. 2017. Transcriptome Association Identifies Regulators of Wheat Spike Architecture. Plant Physiol 175, 746–757.

Wiersma, J.J., Busch, R.H., Fulcher, G.G., and Hareland, G.A. 2001. Recurrent Selection for Kernel Weight in Spring Wheat. Crop Science 41, 999–1005.

Xi, Y., Du, Y.-L., Wang, D., Ren, J.-Y., Luo, W.-Y., Peng, Q., Fang, W.-Y., and Li, F.-M. 2024. Wheat genetic progress in biomass allocation and yield components: A global perspective. Field Crops Research 318, 109617.

Xie, Q., Mayes, S., and Sparkes, D.L. 2015. Carpel size, grain filling, and morphology determine individual grain weight in wheat. Journal of Experimental Botany 66, 6715–6730.

Xie, Q., and Sparkes, D.L. 2021. Dissecting the trade-off of grain number and size in wheat. Planta 254, 3.

Yang, X., Wilkinson, L.G., Aubert, M.K., Houston, K., Shirley, N.J., and Tucker, M.R. 2023. Ovule cell wall composition is a maternal determinant of grain size in barley. New Phytologist 237, 2136–2147.

Yang, Z., Bai, Z., Li, X., Wang, P., Wu, Q., Yang, L., Li, L., and Li, X. 2012. SNP identification and allelic-specific PCR markers development for TaGW2, a gene linked to wheat kernel weight. Theor Appl Genet 125, 1057–1068.

Yang, Z., Van Oosterom, E.J., Jordan, D.R., and Hammer, G.L. 2009. Pre-anthesis ovary development determines genotypic differences in potential kernel weight in sorghum. Journal of Experimental Botany 60, 1399–1408.

Yu, X., Li, B., Wang, L., Chen, X., Wang, W., Wang, Z., and Xiong, F. 2015. Systematic Analysis of Pericarp Starch Accumulation and Degradation during Wheat Caryopsis Development. PLoS One 10, e0138228.

Zadoks, J.C., Chang, T.T., and Konzak, C.F. 1974. A decimal code for the growth stages of cereals. Weed Research 14, 415–421.

Zhai, H., Feng, Z., Du, X., et al. 2018. A novel allele of TaGW2-A1 is located in a finely mapped QTL that increases grain weight but decreases grain number in wheat (Triticum aestivum L.). Theor Appl Genet 131, 539–553.

Zhang, Y., Li, D., Zhang, D., et al. 2018. Analysis of the functions of TaGW2 homoeologs in wheat grain weight and protein content traits. Plant J 94, 857–866.

Zhang, Z., Li, J., Zheng, X., et al. 2024. Ovary morphology determines ovary-to-grain transition process and final grain weight potential in wheat. European Journal of Agronomy 159, 127233.

Zhu, Y., Zhang, X., Xiao, Y., Chu, J., and Dai, Z. 2022. Variation of floret development and grain setting characteristics in winter wheat responses to delayed sowing. Journal of the Science of Food and Agriculture 102, 4892–4908.

