## Supplementary Figures S1-S5 for "*Ta*ExpA6, *VRT-A2* and *Ta*GW2 genes differentially affect grain weight, grain number and yield of wheat through their physiological determinants"

### *Ta*ExpA6, *VRT-A2* and *Ta*GW2 genes differentially affect grain weight, grain number and yield of wheat through their physiological determinants

Lucas Vicentin[1,2] and Daniel F. Calderini[2*]

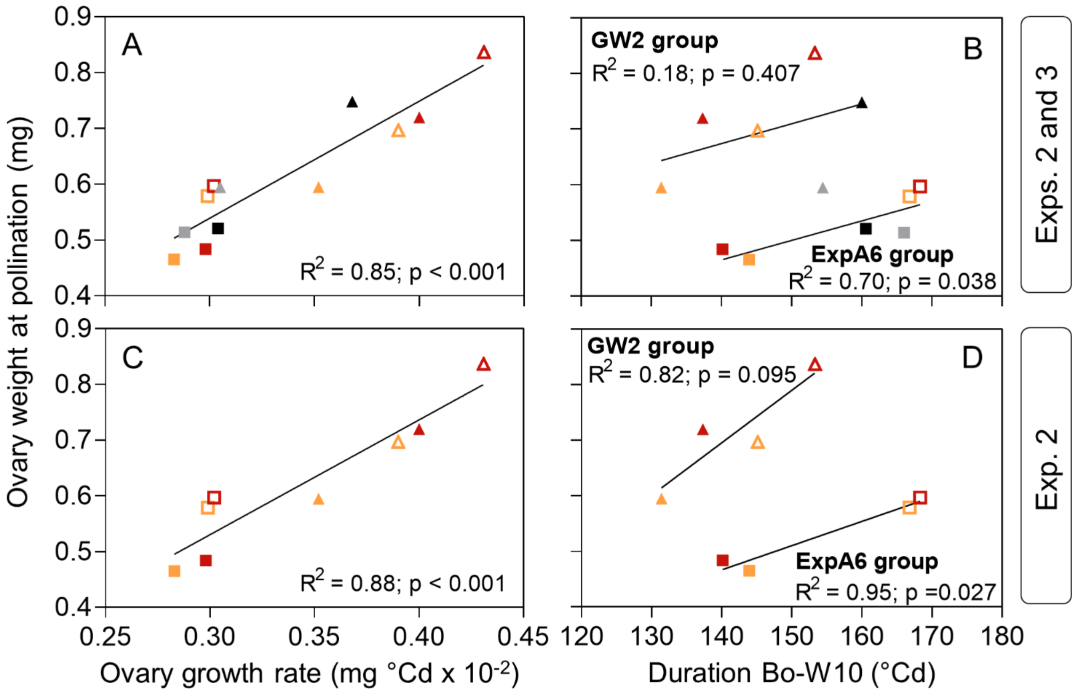


**Figure S1.** Linear relationships between ovary weight at pollination and ovary growth rate in lines ExpA6 (squares) and GW2 (triangles) sown at CPR (closed symbols) and LPR (open symbols) in (A) experiments 2 and 3 and (C) experiment 2 only. Linear relationships between ovary weight at pollination and the duration of the booting – pollination phase in (B) experiments 2 and 3 and (D) experiment 2 only. Data from experiments 2 and 3 are represented, respectively, in red (Mod)/orange (WT), and black (Mod)/grey (WT). Extra sum of squares F test was used to compare the slopes and origins of the regressions between ExpA6 and GW2 groups. When significant differences were found, two equations were fitted.

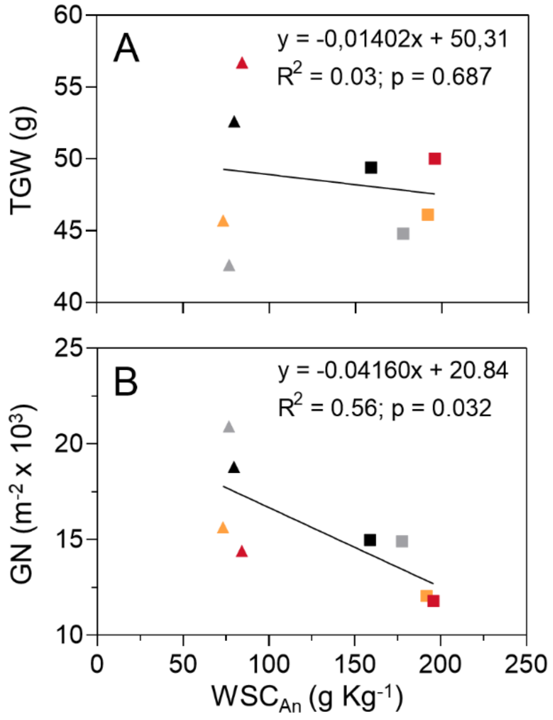


**Figure S2.** Linear relationship between (A) average grain weight and (B) grain number per square metre, and water soluble carbohydrate content in stems at anthesis ($WSC_{An}$) in lines ExpA6 (squares) and GW2 (triangles) sown at CPR plots in experiments 2 and 3. Dark and lighter coloured symbols represent modified (Mod) and WT lines within each group, respectively. Data from experiments 2 and 3 are represented, respectively, in red (Mod)/orange (WT), and black (Mod)/grey (WT).

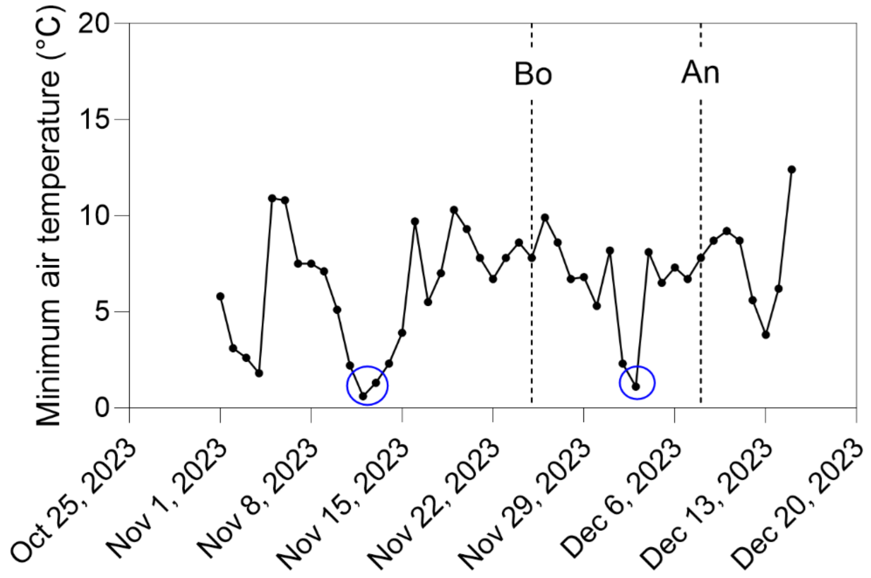


**Figure S3.** Minimum air temperatures below 1.5°C registered in Experiment 3. Temperatures of 0.6, 1.1 and 1.3°C are highlighted with a blue circle. Left and right vertical dashed lines represent booting and anthesis of ExpA6 lines, respectively.

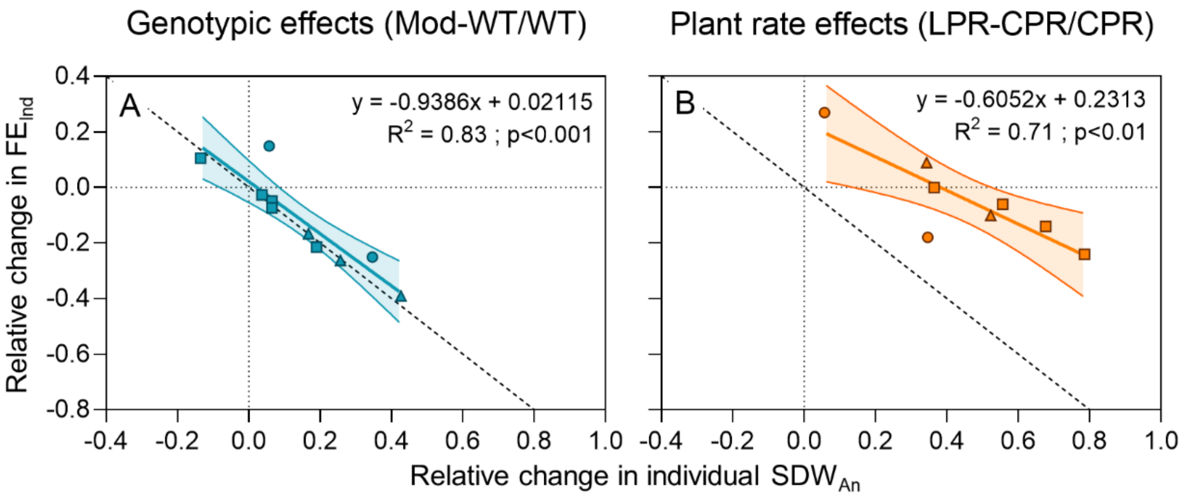


**Figure S4.** Relationship between relative change in individual $SDW_{An}$ and relative change in $FE_{Ind}$ in response to (A) genotypic and (B) plant rate effects in lines ExpA6 (squares), P1xGW2A (circles) and GW2 (triangles). Shaded areas represent 95% confidence bands. Dashed lines represent a slope of -1 (i.e, 1:1 line). Relative change was defined as the ratio between (i) the difference in the trait between the treatment (Modified line or LPR) and the control (WT line or CPR, respectively), and (ii) the trait in the control.

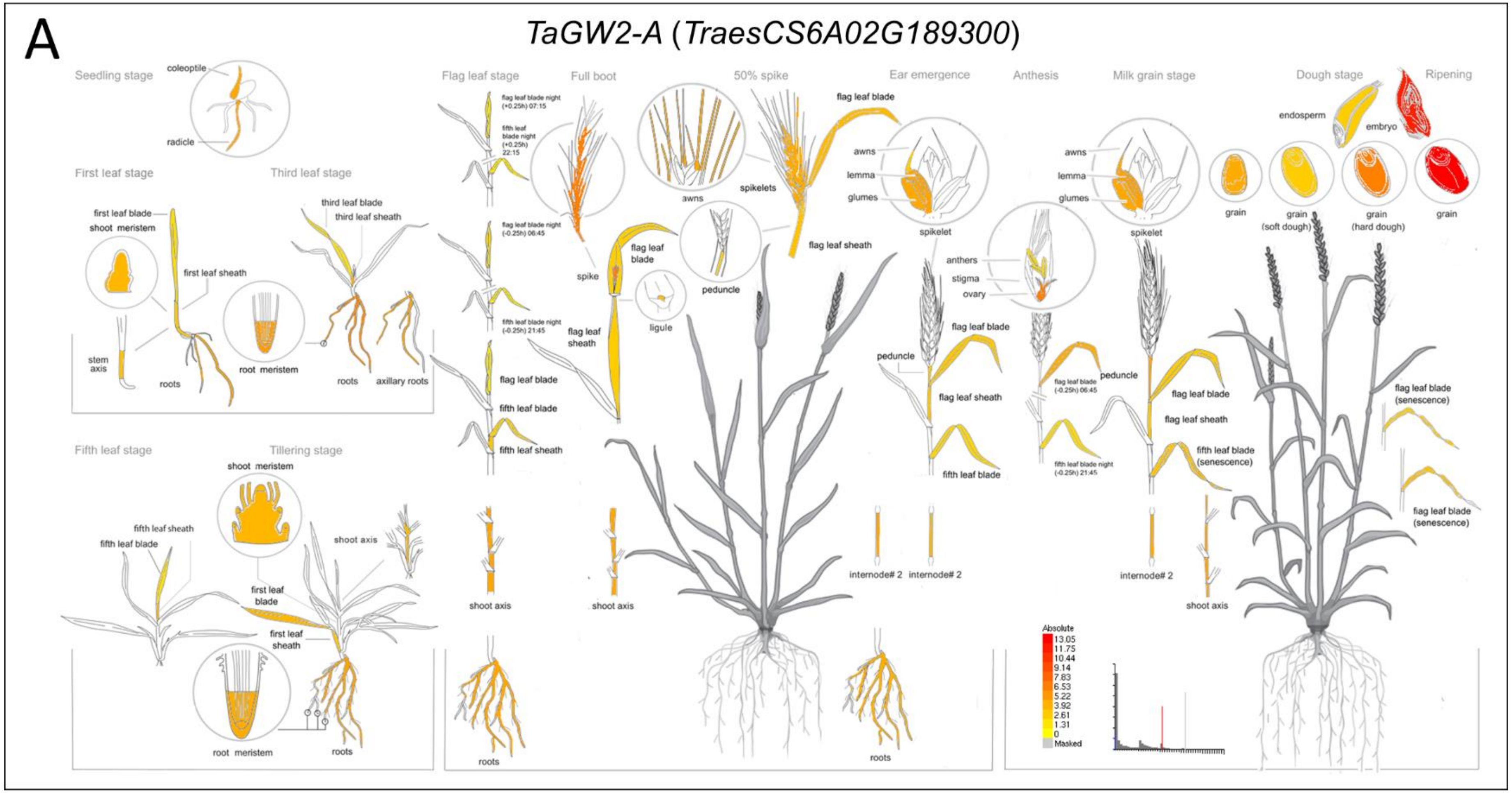


**Figure S5.** Absolute expression values of: (A) TaGW2-A, (B) TaGW2-B and (C) TaGW2-D homeologues in wheat (*T. aestivum*) tissues, as reported by Ramírez-González et al. (2018). Figures retrieved from Wheat eFP Browser (https://bar.utoronto.ca/efp_wheat/cgi-bin/efpWeb.cgi?dataSource=Developmental_Atlas&mode=Absolute). **(Continued)**

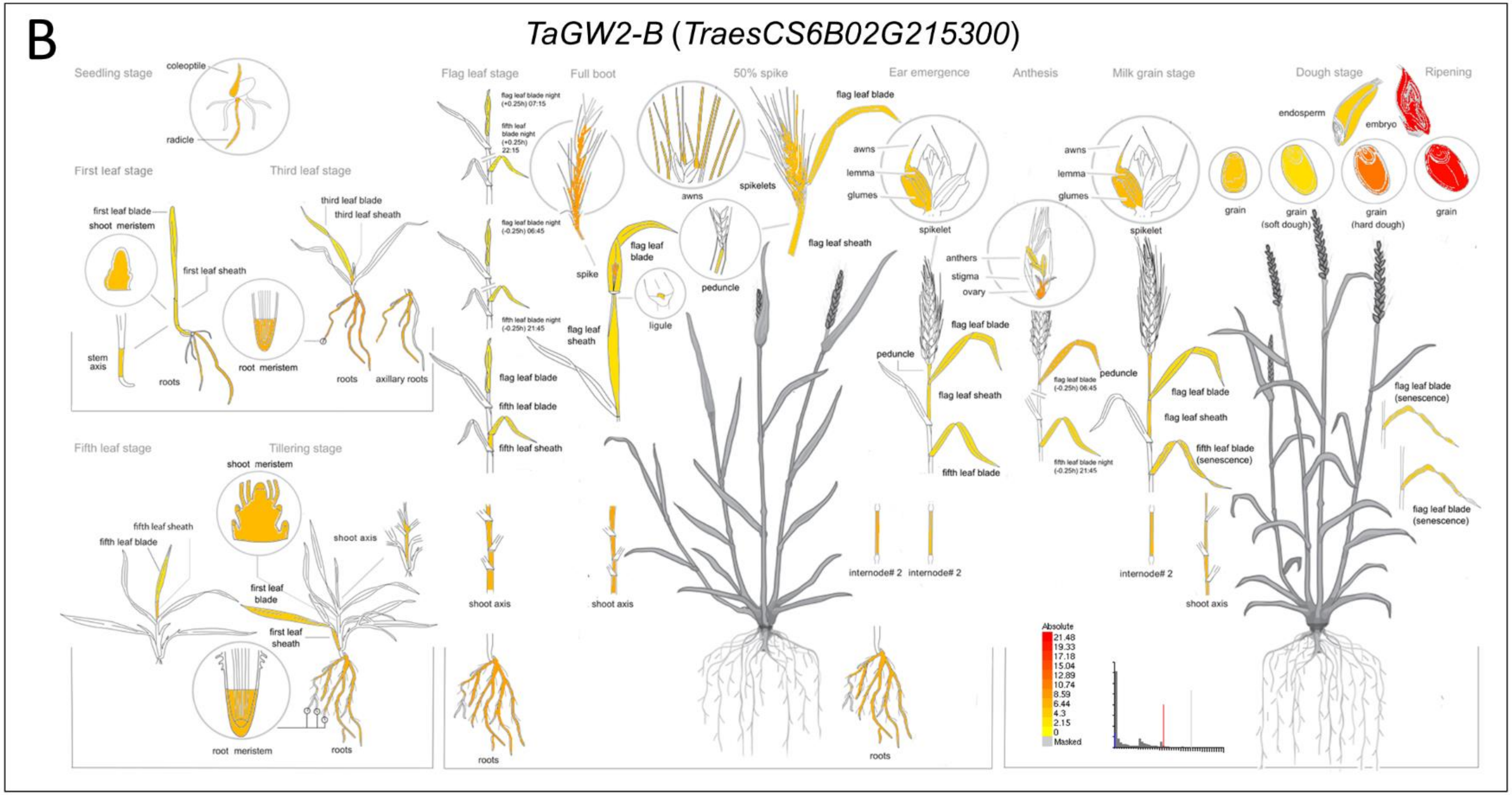


**Figure S5.** Absolute expression values of: (A) TaGW2-A, (B) TaGW2-B and (C) TaGW2-D homeologues in wheat (*T. aestivum*) tissues, as reported by Ramírez-González et al. (2018). Figures retrieved from Wheat eFP Browser (https://bar.utoronto.ca/efp_wheat/cgi-bin/efpWeb.cgi?dataSource=Developmental_Atlas&mode=Absolute). **(Continued)**

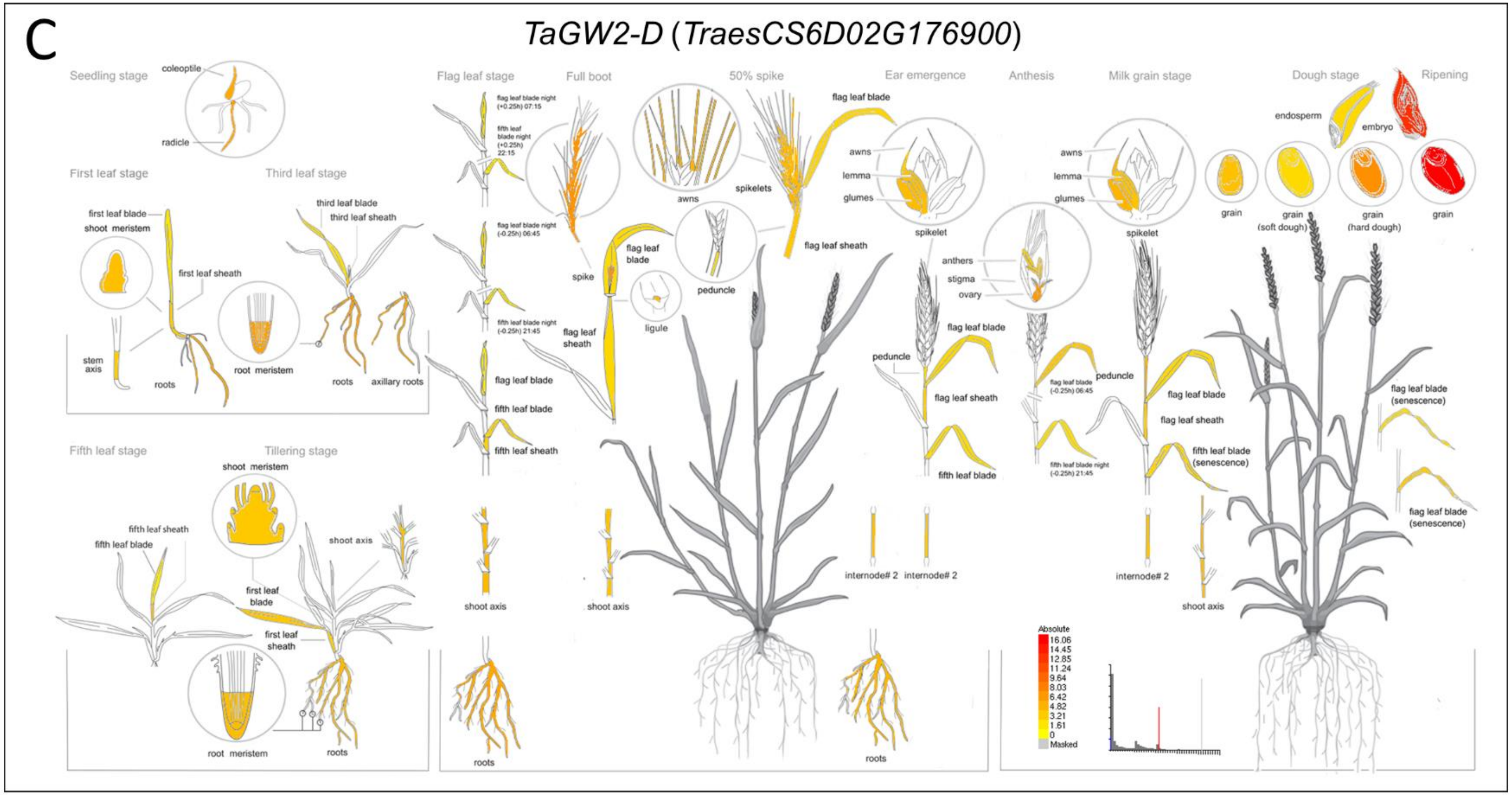


**Figure S5.** Absolute expression values of: (A) TaGW2-A, (B) TaGW2-B and (C) TaGW2-D homeologues in wheat (*T. aestivum*) tissues, as reported by Ramírez-González et al. (2018). Figures retrieved from Wheat eFP Browser (https://bar.utoronto.ca/efp_wheat/cgi-bin/efpWeb.cgi?dataSource=Developmental_Atlas&mode=Absolute).
